# Ancient transposable elements sustain global ecological adaptation despite chronically low nucleotide diversity

**DOI:** 10.64898/2026.03.14.711760

**Authors:** Aidi Zhang, Jordan B. Bemmels, Na Wei

## Abstract

Transposable elements (TEs) are dynamic components of eukaryotic genomes and a major source of structural and regulatory variation, yet their contribution to ecological adaptation over evolutionary time remains unresolved. This uncertainty is particularly acute in species that occupy broad environmental niches despite chronically low nucleotide diversity, where conventional models predict limited adaptive potential. Here we show that ancient TE polymorphisms underpin global ecological adaptation in *Spirodela polyrhiza*, one of the smallest flowering plants with low genome-wide nucleotide diversity. Most TE polymorphisms predate continental population divergence. Cold-season temperature emerges as the dominant selective axis, with adaptive signals overwhelmingly associated with TE polymorphisms rather than SNPs. These adaptive TEs bear signatures of selection on standing variation and are embedded in genomic regions shaped by relaxed purifying selection rather than recent hard sweeps. Our results reveal how ancient TE variation sustains ecological adaptation despite chronically depleted nucleotide diversity, resolving a longstanding evolutionary paradox.

**One sentence summary:** How species adapt with little genetic diversity is a longstanding evolutionary puzzle solved by ancient transposon insertions

## Introduction

Transposable elements (TEs) are pervasive in eukaryotic genomes and represent a uniquely labile source of genomic variation ^1–5^. Through their ability to mobilize, duplicate, and insert across the genome, TEs can generate large-effect mutations and restructure genomic regulatory landscapes in ways that differ fundamentally from point mutations ^2,6–8^. In plants, TE activity is often episodic and environmentally responsive, with bursts of mobilization associated with abiotic and biotic stress, demographic perturbations, and genome-level challenges such as hybridization or polyploidy ^2,4,8,9^. Although most insertions are neutral or deleterious, a subset can be recruited into functional roles, for example by modifying nearby gene expression or donating regulatory or coding sequences, and may persist over long timescales, contributing to standing genomic variation that may shape ecological responses ^2,3,10,11^. These properties have led to the view that TEs contribute not only to genome architecture but also to evolvability, the capacity of genomes to generate adaptive variation under heterogeneous or changing environments ^2–4,6–8,12,13^. Despite the theoretical importance, when and how TE-derived variation contributes to adaptive evolution across natural populations remains poorly understood.

Classic evolutionary theory predicts that adaptive potential scales with standing genomic variation, particularly nucleotide diversity, which fuels selection across heterogeneous environments ^14,15^. Yet this expectation is challenged by species that occupy broad ecological niches despite limited genetic diversity ^10,16^. This pattern is often discussed in the context of the genetic paradox of biological invasions, where rapid adaptation follows severe founder bottlenecks ^16,17^. In such cases, however, reduced diversity is typically transient and subsequently alleviated by demographic recovery, admixture, or repeated introductions ^16^. A more fundamental mystery is whether it is possible for an organism to achieve broad ecological adaptation under chronically low nucleotide diversity. If so, what forms of genomic variation could support it? Addressing this question requires re-evaluating the relationship between nucleotide diversity and adaptive potential in evolutionary theory.

Structural variants, including transposable elements, offer a plausible resolution to this gap. While structural variants such as chromosomal rearrangements, translocations, and copy-number variants have been linked to adaptation ^18–20^, the role of TEs in ecological adaptation has been comparatively underexplored. Unlike single-nucleotide variants, TE polymorphisms can generate large-effect alleles, introduce structural changes, and modify local genomic context, providing forms of variation that may remain accessible to selection even when nucleotide diversity is low ^1,2,4,9,10,16,21^. Importantly, while many TE insertions are deleterious and rapidly purged, others can persist as standing variation, independent of the slow accumulation of point mutations. Empirical studies linking TEs to adaptation have emphasized recent or ongoing transposition, because purifying selection rapidly removes most insertions, leaving detectable signatures primarily among young, low-frequency variants ^11,17,22^. Far less is known about whether ancient, long-standing TE polymorphisms that persist through demographic change and selection can sustain ecological adaptation over evolutionary timescales.

*Spirodela polyrhiza* provides an exceptional natural system for examining how adaptive evolution can proceed under chronically low nucleotide diversity. It is the smallest flowering plant, characterized by rapid clonal growth, a compact genome, and low genome-wide nucleotide diversity worldwide ^23–26^. Yet *Spirodela polyrhiza* is among the most widely distributed angiosperms, occurring on every continent except Antarctica and spanning climatic gradients from cold temperate regions to the tropics ^24^. Such ecological breadth combined with globally low genetic diversity is unusual and appears persistent rather than localized or transient. Previous studies have shown that low diversity in *Spirodela polyrhiza* reflects a low mutation rate and predominantly clonal reproduction, leading to low effective population size and relaxed purifying selection ^23,26^. However, these factors alone do not explain its broad climatic tolerance and global success. The coexistence of long-term genetic paucity with global ecological adaptation therefore presents a fundamental evolutionary paradox. This system provides a natural opportunity to test whether TE variation, if retained over evolutionary time, can contribute to ecological adaptation when conventional genetic diversity is limited.

To address this question, we first reconstruct the demographic history of *Spirodela polyrhiza* to establish the temporal context of its global colonization. We then ask whether TE polymorphisms constitute an important axis of standing genomic variation, and whether this variation is predominantly ancestral or recent. Next, we test the extent to which TE polymorphisms, relative to SNPs, contribute to the climate adaptation across the species’ global range and identify the climatic dimensions most strongly associated with adaptive differentiation. We further assess the genomic context in which climate adaptive TEs reside, asking whether they arise from uniquely unconstrained regions or are selectively filtered from a broadly relaxed genomic background. Finally, we evaluate how TE-dominated variation predicts spatial patterns of climate–genotype mismatch under future climates. Together, these questions address whether ancient TE variation can sustain ecological adaptation under enduring genetic paucity.

## Results

### Recent global divergence and colonization of *Spirodela polyrhiza*

Genome-wide analyses of neutral variation across 245 globally distributed genotypes, each effectively representing population-level sampling owing to predominantly clonal reproduction, revealed that *Spirodela polyrhiza* originated in Asia ^25,26^ and expanded worldwide only recently, during the late Quaternary (Fig. 1a-f). Structure-based clustering identified seven genetic populations across Asia (AS1, AS2), India (IN), Europe (EU1, EU2), and North America (NA1, NA2) (Fig. 1a). This population structure was independently supported by genome-wide principal component analysis of neutral SNPs (Fig. S1) and is consistent with, but refines, earlier continental-scale classifications of Asian, Indian, European, and North American lineages ^25,26^.

**Fig. 1.**
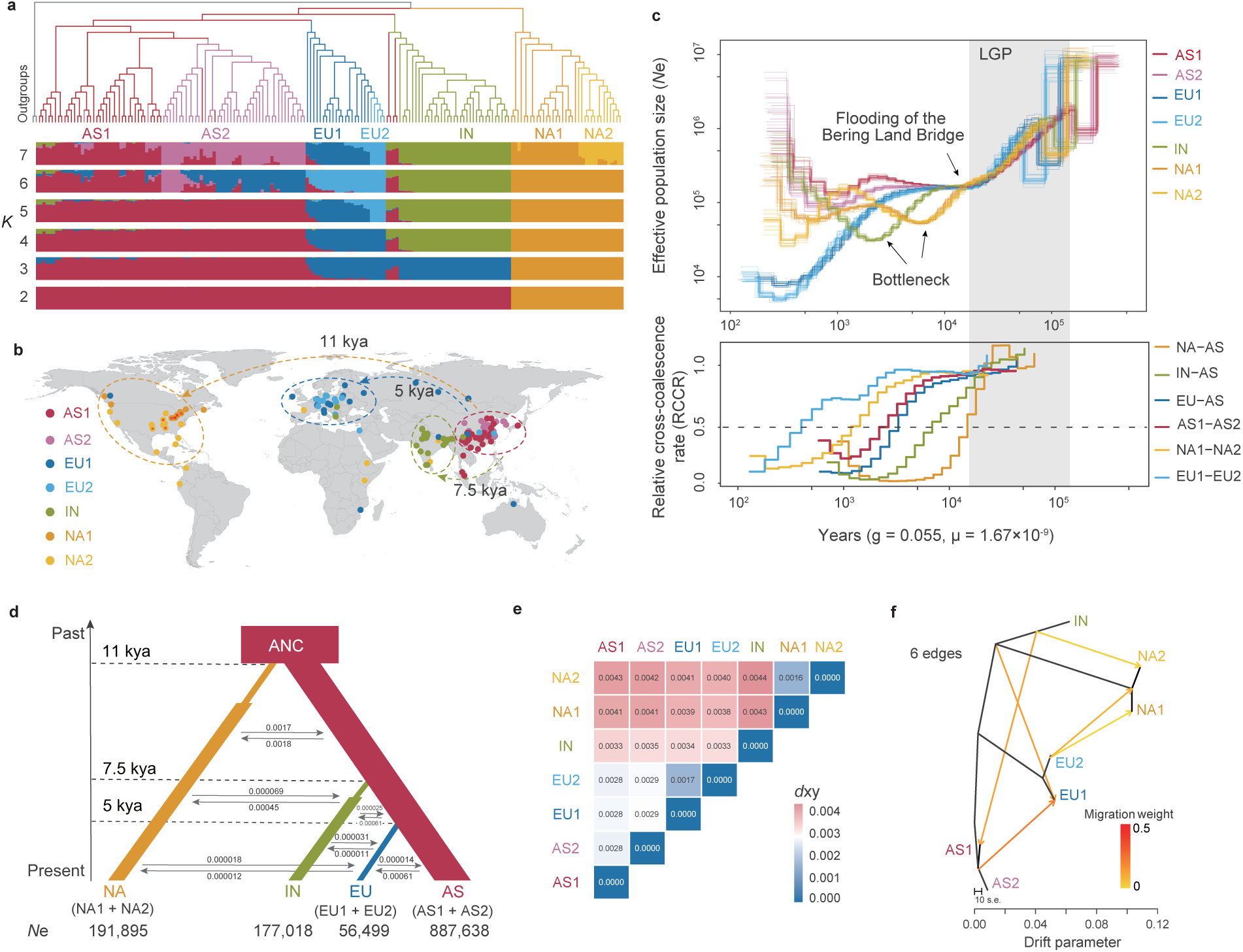
Recent global divergence of *Spirodela polyrhiza* inferred from neutral genomic variation. (a) Maximum likelihood phylogeny of 185 representative accessions (unrelated individuals), rooted with two accessions of *Colocasia esculenta* as outgroups. Colored branches correspond to seven genetic populations identified using ancestry coefficients based on 146,267 putatively neutral 4DTv sites at the optimal clustering level (*K* = 7, determined via cross-validation error), including Asia (AS1, AS2), India (IN), Europe (EU1, EU2), and North America (NA1, NA2). Bar plots show individual ancestry proportions for *K* = 2–7. (b) Geographic distribution and inferred historical dispersal routes. Samples newly sequenced in this study are marked by red ‘×’ (*n* = 17). Dashed ellipses denote regional population clusters and arrows indicate inferred historical migration directions with estimated divergence times (thousand years ago, kya). (c) The dynamics of effective population size (*N*e) and the relative cross-coalescence rate (RCCR) using MSMC2. Shaded region denotes the Last Glacial Period (115-11.7 kya). Major demographic events are indicated, including colonization-associated bottlenecks and the final inundation of the Bering Land Bridge. The dashed horizontal line (RCCR = 0.5) approximates the divergence time between populations. Estimates assumed a sexual generation of 20 days (gen = 0.055) and a mutation rate of 1.67 × 10^−9^ per site per sexual generation. (d) Schematic representation of the best-supported demographic model inferred using fastsimcoal2 (see also Fig. S2). Box widths are proportional to estimated effective population sizes (*N*e). Bottlenecks following lineage splitting in North America and India were included to model dispersal out of Asia. ‘ANC’ represents the common ancestor. Grey arrows denote migration rate per generation. (e) Pairwise absolute nucleotide divergence (*d*xy) estimated in non-overlapping 50-kb windows. Cells are colored by *d*xy values, with red indicating higher divergence and blue indicating lower divergence. (f) Gene flow inferred using TreeMix, showing that global colonization was accompanied by gene flow. Arrows denote migration events, and color intensity represents migration weight.

Demographic modeling supported a sequential Holocene divergence from Asia (Fig. 1b-d; Fig. S2). North American lineages diverged first (∼11 kya), broadly coincident with late-glacial to early-Holocene restructuring of northern dispersal corridors, including the final stages of the Bering Land Bridge. This was followed by divergence of the Indian lineage (∼7.5 kya) during the mid-Holocene monsoon transition, and the most recent split of European lineages (∼5 kya) in the late Holocene. While North American and Indian populations experienced colonization-associated bottlenecks followed by partial recovery, European populations underwent sustained effective population size decline to the present (Fig. 1).

Absolute nucleotide divergence (*d*xy) between populations mirrored this temporal ordering, with progressively greater divergence between Asia and derived continental populations (Fig. 1e). These patterns were consistent with recent divergence from a shared ancestral gene pool rather than long-term isolation. Despite a largely tree-like population history, TreeMix ^27^ analysis revealed evidence of migration among populations (Fig. 1f; Fig. S3), indicating that global colonization was accompanied by gene flow, consistent with secondary contact or multiple introductions rather than strictly unidirectional dispersal.

### Transposable element polymorphisms as an important axis of standing genomic variation

Despite continental-scale sampling, genome-wide nucleotide diversity remains low (π = 0.00073-0.0018; Fig. S1 and Table S1), placing *Spirodela polyrhiza* toward the lower end of π estimates reported across eukaryotes ^28^. Importantly, these continental-scale π estimates primarily reflect divergence among geographically distant locations, as natural populations are predominantly clonal and often exhibit little or no within-population variation ^23,29^. Consistent with this structure, observed heterozygosity was also limited (Fig. S1). Given the low standing SNP variation worldwide ^23,25,26^, we next examined TE polymorphisms as an additional, largely independent source of genomic variation in *Spirodela polyrhiza*.

Transposable elements accounted for approximately 15% of the *Spirodela* genome (*c*. 158 Mb), similar to estimates reported for *Arabidopsis* ^9^. Across the global collection, we identified 14,094 polymorphic TE insertions that were broadly distributed along chromosomes (Fig. 2a and Fig. S4). For comparison, we detected 1,675,572 genome-wide SNPs (Fig. S4), consistent with previous global surveys of *Spirodela polyrhiza* ^25^, of which 232,169 had a minor allele frequency (MAF) ≥ 0.05. Thus, TE polymorphisms represent an important and distinct component of genomic variation in *Spirodela polyrhiza*. These polymorphic TEs were dominated by LTR retrotransposons and DNA transposons (Fig. S5), consistent with patterns reported across plant genomes ^4,9,11^. In contrast to SNPs, polymorphic TEs showed a higher relative representation in exonic regions, whereas SNPs occurred more frequently in intergenic and intronic regions (Fig. 2a, inset; Table S2). This structural distinction indicates that TE polymorphisms are more likely than SNPs to intersect coding sequences ^22,30^, with potential consequences for gene function.

**Fig. 2.**
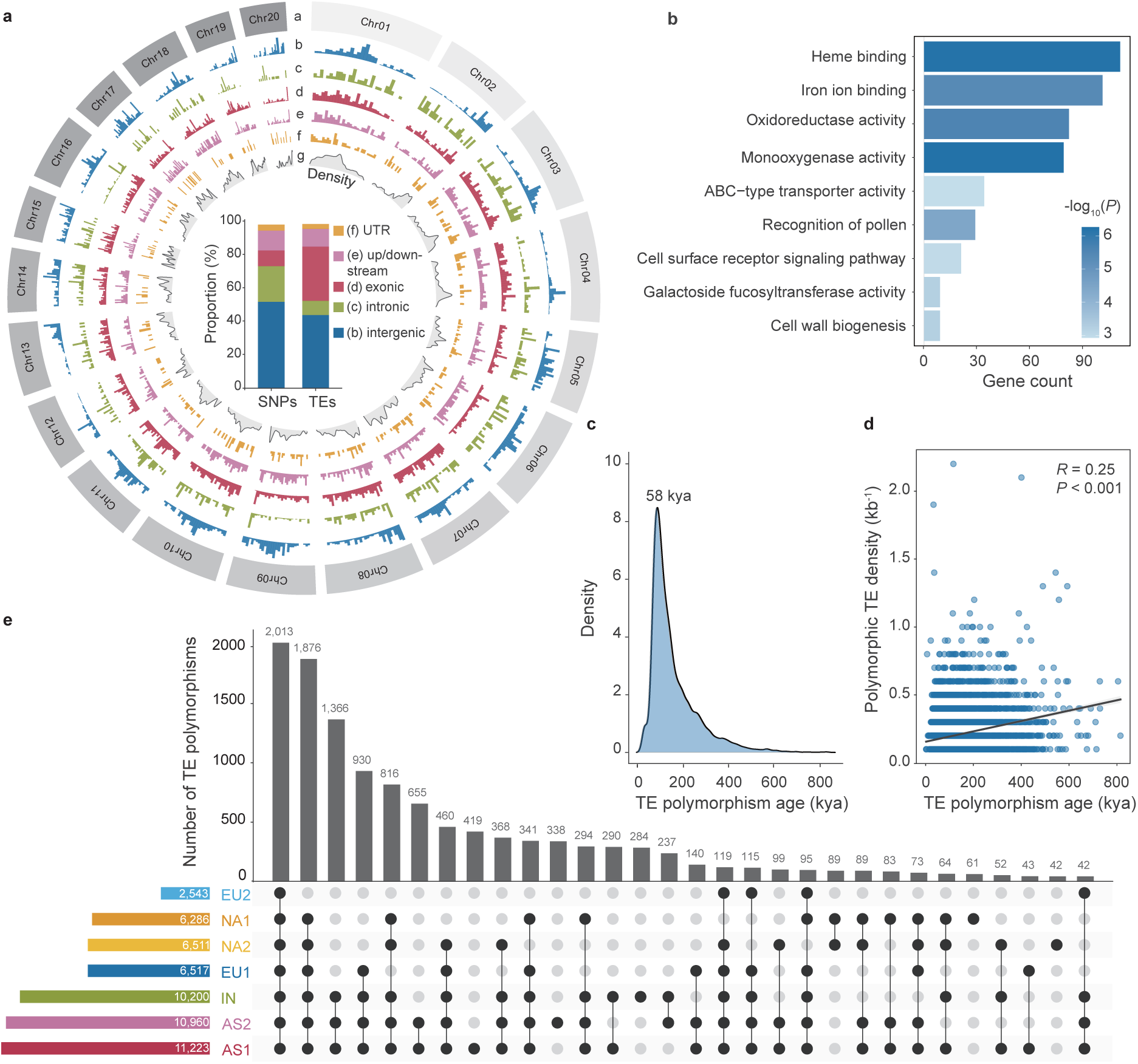
TE variation constitutes an important axis of standing genomic variation in *Spirodela polyrhiza.* (a) Genome-wide distribution of 14,094 polymorphic TEs identified in the global collection. Tracks (outer to inner) represent chromosomes, intergenic, intronic, exonic, upstream/downstream (±2 kb), and UTR regions; the innermost track shows polymorphic TE density (number per 10-kb non-overlapping window). The inset histogram compares the proportional distribution of SNPs and polymorphic TEs across genomic regions. (b) Gene Ontology (GO) enrichment of genes harboring polymorphic TEs (Benjamini–Hochberg adjusted *P* < 0.05). Bar length indicates gene count; color intensity corresponds to -log10 (adjusted *P*). (c) Distribution of estimated ages of polymorphic TEs (kya), with a peak around ∼58 kya. (d) Positive correlation between polymorphic TE density and mean TE age across non-overlapping 10-kb windows (*n* = 6,922) based on Pearson’s correlation coefficient (*R*), supporting the long-term retention of ancient TE variation. (e) Distribution of polymorphic TEs across the the seven genetic populations. Horizontal colored bars indicate the total number of polymorphic TEs detected per population. Vertical bars represent shared and population-specific polymorphic TEs.

Genes harboring polymorphic TEs were significantly enriched for functions related to redox processes, metal ion binding, and membrane transport, including heme binding, iron ion binding, oxidoreductase activity, monooxygenase activity, and ABC-type transporters (Fig. 2b). Enrichment was also observed for receptor-mediated signaling and cell surface-associated processes. These enrichments indicate that polymorphic TEs preferentially intersect genes involved in redox regulation, transport, and environmental sensing, processes we hypothesize to be essential for coping with fluctuating aquatic environments.

Polymorphic TEs were predominantly ancient, with a pronounced age peak around ∼58 kya (Fig. 2c), substantially predating the global population divergences of *Spirodela polyrhiza* (∼5-11 kya; Fig. 1). This peak coincided with the last glacial period (Fig. 1c) and has been seen in other species as well ^22,30^, indicating that most standing TE polymorphisms arose prior to global colonization and persisted through subsequent demographic bottlenecks. Consistent with this pattern, polymorphic TE density increased with inferred TE age (Fig. 2d), supporting long-term retention of ancient TE variation.

An UpSet analysis revealed extensive sharing of TE polymorphisms across populations, with most variation shared across multiple populations (Fig. 2e). Asian populations (AS1 and AS2) harbored the largest pools of TE polymorphisms and contributed disproportionately to shared variation, consistent with their role as ancestral reservoirs. Anchoring TE ages to inferred population divergence times (Fig. 1) identified very few lineage-specific, post-divergence TE polymorphisms: none in North American or European populations and only two in India. Together, these results indicate that contemporary TE variation across *Spirodela polyrhiza* populations overwhelmingly reflects ancestral polymorphisms that predated global divergence.

### Climate-associated TE polymorphisms dominated putatively adaptive signals

Given that TE polymorphisms constitute abundant standing genomic variation, we next asked the extent to which they, relative to SNPs, contribute to the climate adaptation of *Spirodela polyrhiza* across its global range. Multivariate genotype–environment association (using RDA) ^31^ yielded concordant rankings of climatic predictors for TE polymorphisms (*n* = 14,094) and SNPs (*n* = 232,169), with minimum temperature of the coldest month (Bio06) consistently emerging as the strongest axis of association (Fig. 3a), indicating cold-season temperature as a dominant climatic constraint.

**Fig. 3.**
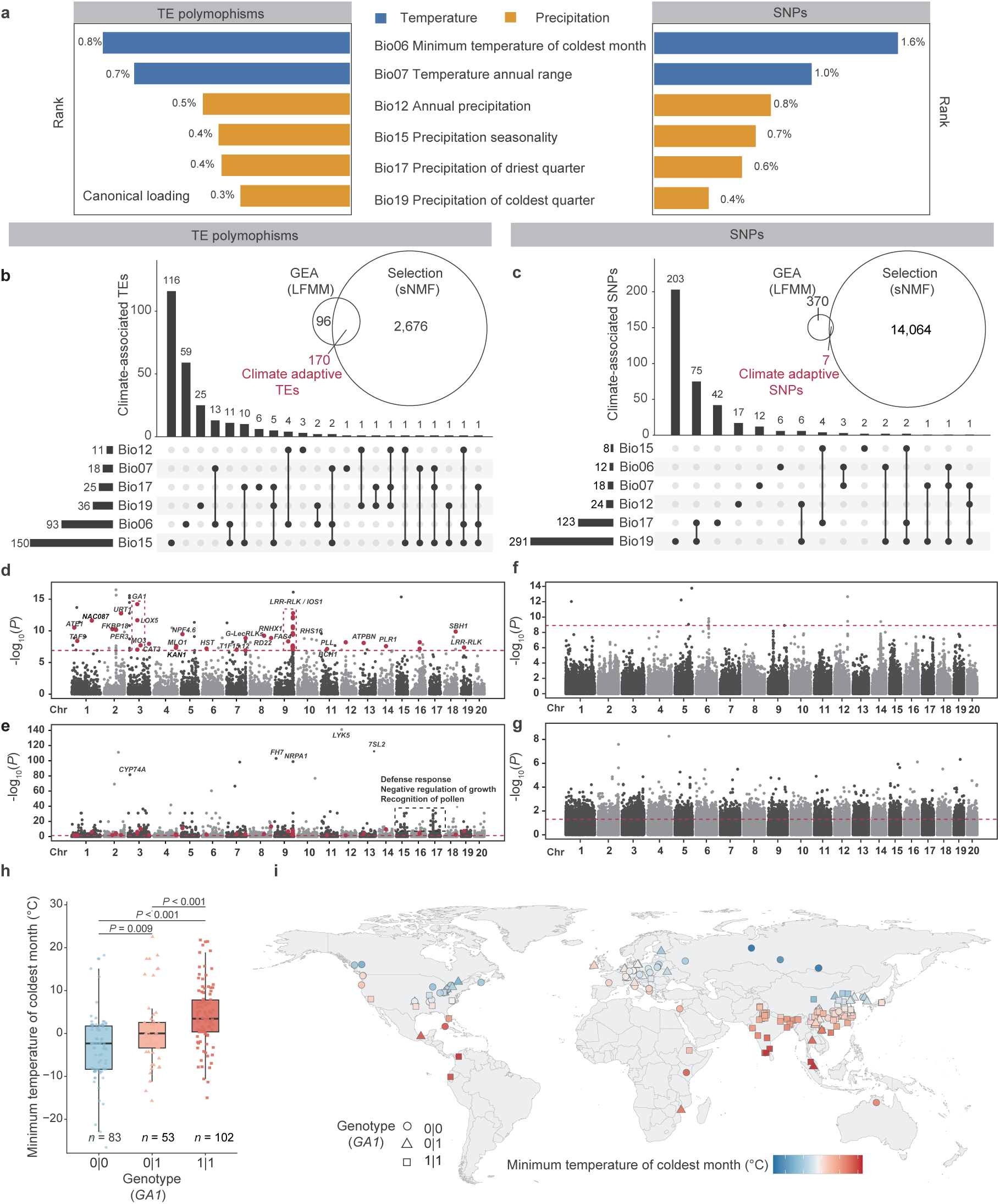
Climate-associated TE variation dominates adaptive signals across the global range of *Spirodela polyrhiza*. (a) Multivariate genotype-environment association (GEA) using redundancy analysis (RDA) for TE polymorphisms (*n* = 14,094) and SNPs (*n* = 232,169; MAF ≥ 0.05). Both methods yielded concordant rankings of bioclimatic predictors, with Minimum Temperature of Coldest Month (Bio06) emerging as the strongest axis of association. Bars represent canonical loadings of the six bioclimatic variables ranked by contribution. Blue indicates temperature-related variables and orange indicates precipitation-related variables. (b, c) Identification of climate adaptive TE polymorphisms (b) and SNPs (c) across six bioclimatic variables. Climate adaptive loci were defined as the intersection between climate-associated loci identified by GEA (latent factor mixed model, LFMM) and loci under selection inferred using sparse non-negative matrix factorization algorithms (sNMF). Numbers indicate loci identified by each approach and their overlap. UpSet plots show the distribution of climate-associated loci across single and multiple bioclimatic variables. (d, e) Manhattan plots of climate adaptive TEs associated with Minimum Temperature of Coldest Month (Bio06). LFMM-based GEA signals are shown in (d), with the dashed horizontal line indicating the empirical threshold derived from null models generated by randomizing geographic coordinates to disrupt genotype-environment associations. sNMF-based selection signals are shown in (e), with the dashed horizontal line indicating the empirical threshold derived from putatively neutral 4DTv sites. Climate adaptive loci supported by both analyses are highlighted in red. (f, g) Manhattan plots of climate adaptive SNPs associated with Bio06. LFMM-based GEA signals are shown in (f), and sNMF-based selection signals are shown in (g). No loci were jointly supported by both analyses. (h) Boxplots of minimum temperature of the coldest month (°C) among the three *GA1* genotypes identified as climate adaptive in (d). *GA*1 encodes a key enzyme in gibberellin biosynthesis. Boxes represent interquartile ranges, center lines indicate medians, and whiskers extend to 1.5× the interquartile range. Pairwise differences were assessed using two-sided Wilcoxon rank-sum tests. (i) Geographic distribution of the three *GA1* genotypes shown in (h). Symbols denote genotypes and are colored according to the minimum temperature of the coldest month (°C).

As expected for largely ancestral variants, TE polymorphisms also captured strong population structure concordant with neutral SNPs (Fig. S5), highlighting the need to account for demography when identifying adaptive signals. Using LFMM ^32^ with demography explicitly considered, we identified 266 climate-associated TE polymorphisms across six climatic variables (Fig. 3b; Figs. S6-S10). To distinguish associations driven by selection from those reflecting other evolutionary processes, including genetic drift, we further tested TE polymorphisms for signatures of selection (Fig. 3b). A subset of 170 TE polymorphisms overlapped between climate association and selection signals and were defined as climate adaptive TEs (Fig. 3b; Table S3).

Independent validation using BayeScan ^33^ supported this distinction: climate adaptive TEs showed signatures of positive selection, whereas non-adaptive TEs were largely neutral (Fig. S11). Genomic annotation revealed that more than half (54%) of the climate adaptive TEs were gene-associated (exonic and regulatory regions) and 37% were located in regulatory space, with the remainder intergenic (Table S3).

In contrast, applying the same framework to SNPs identified only seven climate adaptive SNPs (Fig. 3c). Although many SNPs were detected as climate-associated (*n* = 370) and numerous exhibited signatures of selection (*n* = 14,064), the overlap between these categories was minimal. This limited concordance suggests that climate-associated SNPs rarely coincide with loci exhibiting strong selection signals. Such decoupling may reflect several non-mutually exclusive processes, including diffuse polygenic responses, background or linked selection, or selection driven by non-climatic factors. By contrast, TE polymorphisms more frequently exhibited coordinated signals of climate association and positive selection.

Focusing on cold-season temperature (Bio06), several climate adaptive TEs mapped near genes involved in growth and stress signaling, including a GA1 homolog, a key enzyme in gibberellin biosynthesis that links temperature cues to growth regulation ^34–36^, and an LRR receptor-like kinase annotated as IOS1, implicated in stress signaling ^37^ (Fig. 3d). Additional prominent TE selection peaks lacked Bio06 association and were linked to genes involved in defense responses, growth regulation, and pollen recognition, suggesting selective pressures beyond cold-season temperature (Fig. 3e). In contrast, SNP-based scans revealed no climate adaptive candidates for Bio06 (Fig. 3f,g). Consistent with climate association, TE genotypes at the GA1-linked locus exhibited a stepwise relationship with Bio06 across the global temperature gradient (Fig. 3h,i).

### Adaptive TE windows showed signatures of selection on standing variation

To examine the genomic context in which climate adaptive TEs reside, we compared SNP-based population genetic statistics between genomic windows harboring climate adaptive TEs and those containing non-adaptive TEs. Contrary to expectations under classic hard selective sweeps, adaptive TE windows exhibited significantly higher nucleotide diversity (π) than non-adaptive TE windows using a resampling approach (Fig. 4a,b; see Methods).

**Fig. 4.**
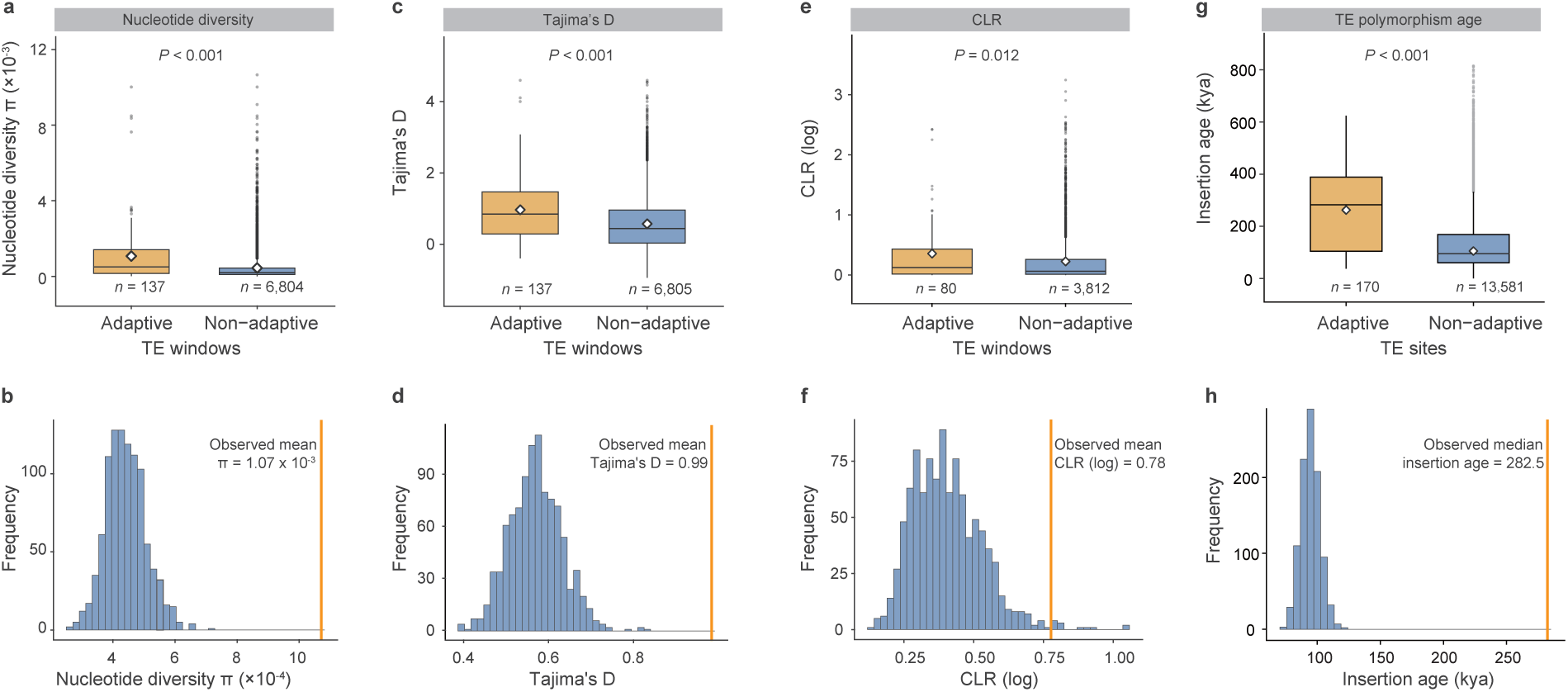
Climate adaptive TE windows show signatures of selection on standing variation and deep evolutionary origins. (a, b) Nucleotide diversity (π) calculated from SNPs within non-overlapping 10-kb windows containing climate adaptive TEs (adaptive TE windows) and windows containing non-adaptive TEs (non-adaptive TE windows). Because non-adaptive windows outnumbered adaptive windows (a), non-adaptive windows were randomly subsampled 1,000 times to generate a permutation-based null distribution (blue histograms in b). Statistical significance was assessed using a two-sided permutation test (see Methods). The orange vertical line in (b) indicates the observed mean π of adaptive windows. (c, d) Tajima’s D calculated from SNPs was compared between adaptive and non-adaptive windows. (e, f) Selective sweep signals were assessed from SNPs using SweeD, and the composite likelihood ratio (CLR) was compared between adaptive and non-adaptive windows. (g, h) Climate adaptive TE loci were much older than non-adaptive TE loci. Across panels, boxplots show medians (center lines), interquartile ranges (boxes), and whiskers extending to 1.5× the interquartile range; diamonds denote means. Window numbers are indicated in (a-f) and TE numbers are indicated in (g-h).

Consistent with elevated diversity, Tajima’s D was also higher in adaptive TE windows (Fig. 4c,d), indicating an excess of intermediate-frequency variants that may reflect spatially heterogeneous selection in the global collection. These patterns are inconsistent with recent hard sweeps, but instead align more closely with scenarios involving soft sweeps or polygenic adaptation, or potentially relaxed purifying selection (which was ruled out later in the analysis). Supporting this interpretation, adaptive TE windows showed elevated composite likelihood ratio (CLR) ^38^ values relative to non-adaptive windows (Fig. 4e,f), further indicating the presence of detectable selection signals.

Climate adaptive TEs were significantly older than non-adaptive TEs (Fig. 4g,h), indicating that adaptive TE polymorphisms predominantly derived from long-standing ancestral insertions rather than recent, post-divergence activities. Together, these results suggest that climate adaptation in *Spirodela polyrhiza* is associated with ancient TE polymorphisms residing in genomic regions shaped by selection on standing variation, consistent with soft sweeps or polygenic adaptation, rather than by recent hard selective sweeps that typically reduce surrounding diversity.

### Relaxed purifying selection shaped the genomic background of TE polymorphisms

Because the efficacy of purifying selection determines which variants persist as standing variation available for adaptation, we next examined how selective constraint is distributed across the *Spirodela* genome and whether it is associated with TE-rich regions.

Genome-wide estimates of π_N_/π_S_ revealed pervasive purifying selection in *Spirodela polyrhiza* (median π_N_/π_S_ < 1), accompanied by substantial heterogeneity, including a long tail of genomic windows with elevated π_N_/π_S_ indicative of locally relaxed constraint (Fig. 5a). Windows with elevated π_N_/π_S_ were enriched for genes associated with negative regulation of growth (Fig. 5b,c), indicating relaxed purifying selection on growth-inhibitory pathways. In a rapidly proliferating aquatic plant such as *Spirodela polyrhiza*, reduced constraint on growth repression is consistent with selection favoring rapid vegetative expansion. Such relaxed constraint may also render growth regulatory genes permissive targets for adaptive regulatory modification by transposable elements (Fig. 3e).

**Fig. 5.**
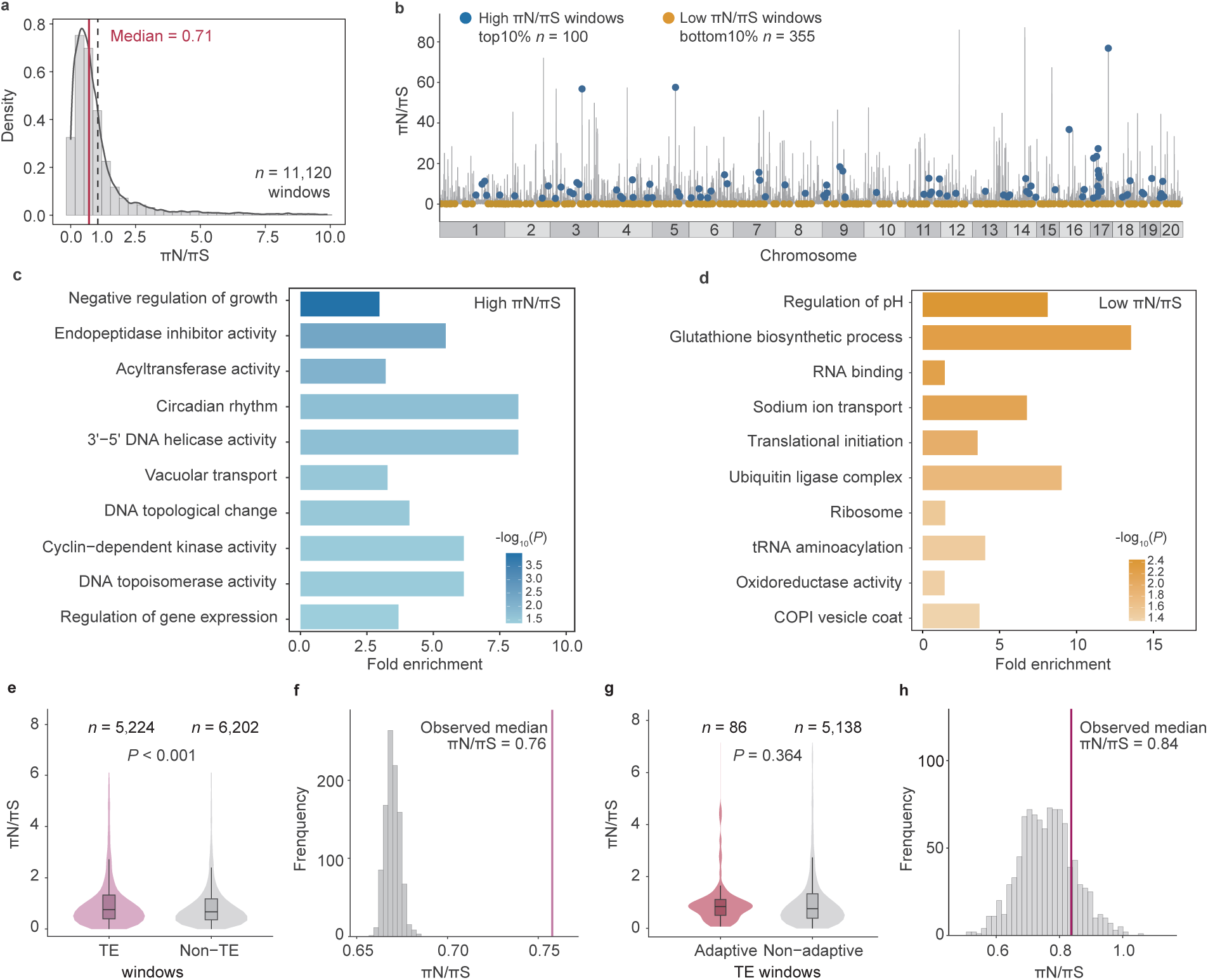
Relaxed purifying selection shapes the genomic background of TE variation. (a) Distribution of the ratio of nonsynonymous to synonymous nucleotide diversity (πN/πS) across non-overlapping 10-kb genomic windows (*n* = 11,120). The red vertical line indicates the median (0.71). (b) Genome-wide distribution of πN/πS across chromosomes. Each line represents a 10-kb window. Windows were ranked by πN/πS values: the top 10% (high πN/πS) are shown in blue and the bottom 10% (low πN/πS) are shown in orange for downstream Gene Ontology (GO) enrichment analysis. (c, d) GO enrichment analysis of genes located in windows with the top 10% (c) and the bottom 10% of πN/πS. Bar length indicates fold enrichment and color intensity corresponds to −log10 (adjusted *P*). (e, f) Comparisons of πN/πS between 10-kb windows containing TEs (TE windows, *n* = 5,224) and windows lacking TEs (non-TE windows, *n* = 6,202). Violin plots with embedded boxplots are shown; center lines indicate medians and whiskers extend to 1.5× the interquartile range. Because non-TE windows outnumbered TE windows, non-TE windows were randomly subsampled 1,000 times to generate a permutation-based null distribution (grey histograms in f). Statistical significance was assessed using a two-sided permutation test (see Methods). The pink vertical line in (f) indicates the observed median πN/πS of TE windows. (g, h) Within TE windows (*n* = 5,224), πN/πS was compared between windows containing climate adaptive TEs (*n* = 86) and windows containing non-adaptive TEs (*n* = 5,138). Statistical significance was assessed using a two-sided permutation test.

In contrast, windows with low πN/πS were enriched for pathways underlying cellular homeostasis, including regulation of pH and glutathione biosynthesis (Fig. 5d). These functions are essential for maintaining ion and redox balance under fluctuating aquatic conditions ^39^, consistent with strong purifying selection acting on core physiological processes in *Spirodela polyrhiza*.

TE-containing windows exhibited significantly higher πN/πS than non-TE windows (Fig. 5e,f), indicating that TE polymorphisms were preferentially retained in genomic regions under weaker purifying selection. In contrast, windows harboring climate adaptive TEs did not differ in πN/πS from those containing non-adaptive TEs (Fig. 5g,h), indicating that climate adaptive TEs were not associated with unusually weak constraint relative to other TE-containing regions, but instead represented a subset selectively favored from a broadly relaxed genomic background.

### Genetic offset captures spatial heterogeneity in climate mismatch

To assess how climate adaptive TE polymorphisms translate into spatial patterns of future climate–genotype mismatch, we quantified genetic offset with respect to cold-season temperature (Bio06) using two complementary approaches: LFMM-based genetic gap ^32^ and Gradient Forest ^40^ (Fig. 6). Despite differences in absolute magnitude, both methods revealed highly consistent patterns (Fig. 6a,c), with North American and European populations, especially NA1 and EU1, exhibiting the highest offset. Geographic projections further revealed pronounced spatial heterogeneity in genetic offset, largely structured along latitudinal gradients (Fig. 6b,d). Together, these results show that TE-dominated climate adaptation gives rise to spatially heterogeneous genetic mismatches to future climatic change.

**Fig. 6.**
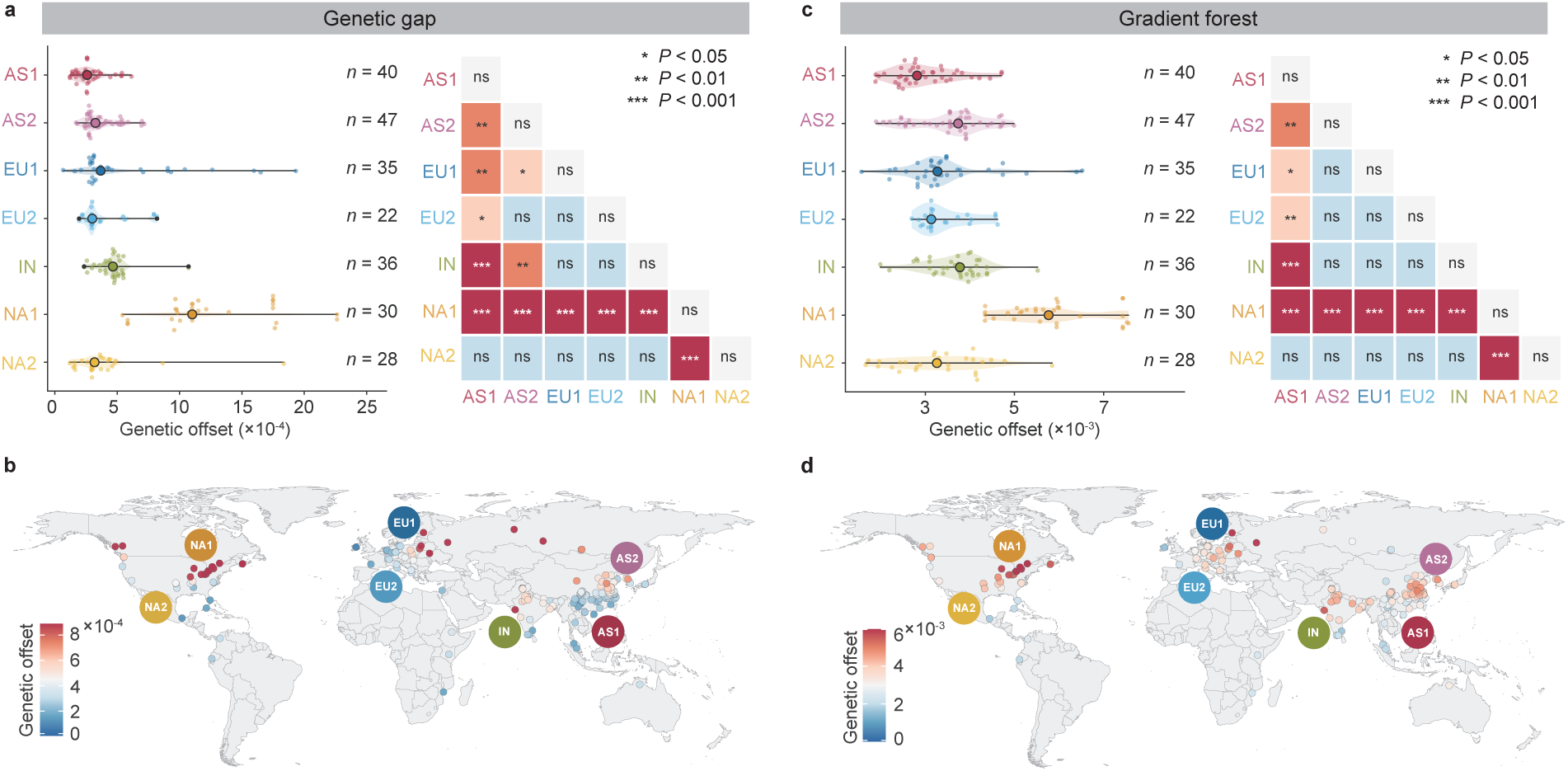
TE variation predicts genetic offset under projected future climates. (a-b) Genetic offset estimated from climate adaptive TEs associated with the Minimum Temperature of Coldest Month (Bio06), using a latent factor mixed model (LFMM) based genetic gap approach. Future climate projections correspond to 2081-2100 under SSP5-8.5 scenario. In (a), violin plots with quasirandom jittered points represent individual genetic offset values. Filled circles indicate medians, and horizontal lines denote the full range (minimum to maximum) within each population. Heatmaps summarize pairwise comparisons (two-sided Wilcoxon rank-sum test with Benjamini-Hochberg correction; *** *P* < 0.001, ** *P* < 0.01, * *P* < 0.05). In (b), colors represent estimated genetic offset values, with red indicating higher offset and blue indicating lower offset. Population names are indicated with colored circles. (c, d) Genetic offset estimated from climate adaptive TEs associated with Bio06 using Gradient Forest.

## Discussion

Our results reveal that global ecological adaptation can be sustained despite limited standing SNP variation. In *Spirodela polyrhiza*, climate-associated adaptation is dominated by TE polymorphisms rather than SNPs and unfolds against a background of low genome-wide nucleotide diversity across the species’ worldwide distribution. This finding challenges the prevailing assumption that adaptive potential necessarily scales with nucleotide diversity and recombination, and instead identifies ancient TE polymorphisms as a durable substrate for adaptative evolution. Notably, climate adaptive TEs substantially predate recent continental divergence, remaining available to selection over hundreds of thousands of years across fluctuating environments. Together, these results demonstrate that ancient TE variation can decouple global ecological breadth from genome-wide nucleotide diversity, revealing an alternative genetic foundation for long-term adaptive potential.

The evolutionary dynamics revealed here are distinct from those typically invoked to explain rapid adaptation under low genetic diversity in biological invasions. In invasive systems, reduced diversity usually reflects recent founder events and is often transient, with adaptive potential restored through demographic expansion, admixture, or repeated introductions ^16,41^.

Transposable elements have been implicated in such contexts primarily through recent or ongoing transposition that generates novel variants during episodes of demographic or environmental disruption ^16,17,42^. By contrast, *Spirodela polyrhiza* exhibits persistently low nucleotide diversity across its global range (Fig. S1), with no evidence for post-bottleneck genetic recovery or continual mutational replenishment ^23,25,26^. Its ecological success therefore cannot be attributed to invasion-associated dynamics or to recent bursts of transposition. Instead, the adaptive signal uncovered here is dominated by ancient TE polymorphisms that have persisted through multiple demographic transitions (Fig. 3 and Fig. 4). This distinction is crucial, indicating a mode of adaptation that is mechanistically and temporally distinct from invasion genetics.

The dominance of ancient TE polymorphisms in climate adaptation raises a central question: where did this variation originate, and why has it persisted for so long? In *Spirodela polyrhiza*, the age distribution of polymorphic TEs shows a pronounced peak around the last glacial period (Fig. 2), substantially predating global colonization (Fig. 1). Similar late-Pleistocene pulses of TE insertions have been reported in other plant lineages ^22^, suggesting that glacial–interglacial environmental instability broadly promoted episodic transposition rather than lineage-specific events. Notably, the subset of TEs implicated in climate adaptation is even older (Fig. 4), indicating that adaptive signals draw disproportionately from deeply rooted insertions. Comparative analyses across hundreds of plant genomes further show that the TE families contributing to standing variation in *Spirodela* (Fig. 2 and Fig. S5) are typical of angiosperms ^2,4,5,9,11^, rather than representing an idiosyncratic expansion. Importantly, these ancient insertions were not eliminated during subsequent demographic contractions or range shifts, implying that a subset of TE-derived variants can evade purifying selection over extended evolutionary timescales. Together, these patterns support a view of transposable elements as durable components of genome architecture that can retain adaptive relevance long after periods of elevated activity have passed.

The persistence and adaptive relevance of ancient TE polymorphisms in *Spirodela polyrhiza* likely reflect a distinctive genomic and epigenetic context shaped by clonal reproduction and genome streamlining. Recent work shows that *Spirodela polyrhiza* exhibits a reconfigured TE silencing landscape, characterized by reduced RNA-directed DNA methylation (RdDM) and diminished 24-nt siRNA pathways, with preferential silencing of the relatively few intact TEs, whereas a majority of TEs are ancient and largely degenerated with reduced canonical methylation ^25,43–46^. This architecture plausibly limits new transposition while allowing long-standing TE variants to persist as standing genomic features. In a genome characterized by low recombination, low mutation rate, and chronically low effective population size ^25,26^, newly arising TE insertions are likely to be predominantly deleterious and epigenetically suppressed, whereas ancient TE polymorphisms that have already survived prolonged selective filtering remain available for adaptive sorting. This framework reconciles studies emphasizing TE load ^47,48^ with those implicating TEs in adaptation ^2,22^, including the present study. In *Spirodela polyrhiza*, climate adaptive TEs are not associated with unusually weak constraint but instead represent a selectively favored subset drawn from a broadly permissive genomic background (Fig. 5), highlighting how the evolutionary consequences of transposable elements depend critically on genomic context and life history.

Rather than acting randomly across the genome, climate adaptive TEs preferentially intersect growth-regulatory pathways (Fig. 3), pointing to growth modulation as a target of selection. In a clonally reproducing plant, fitness is tightly coupled to vegetative expansion and seasonal timing, making partial relaxation of growth repression adaptive rather than deleterious. Pathways such as gibberellin biosynthesis integrate temperature cues with growth decisions, providing natural leverage points for regulatory fine-tuning without compromising core cellular functions ^49–51^. In this context, relaxed purifying selection does not indicate genomic decay but reflects an evolutionary strategy that preserves flexibility in growth control while maintaining strong constraint on essential homeostatic processes. We hypothesize that this functional bias helps explain why TE insertions in genic regions can contribute to adaptation without incurring the genetic load typically assumed for coding variation.

Cold-season temperature emerges as a biologically meaningful selective axis for *Spirodela polyrhiza* (Fig. 3). Although the species spans a broad latitudinal range, active growth typically does not take place below ∼8 °C and fronds tolerate temperatures only down to, but not substantially below, the freezing point ^52^. When projected into future climates, TE-dominated adaptation translates into pronounced spatial heterogeneity in genetic offset (Fig. 6), particularly at higher latitudes, indicating uneven capacity to track thermal change. Unlike systems in which climate change may promote adaptive novelty through new TE insertions, adaptation in *Spirodela* appears to rely primarily on ancient standing TE variation, with limited scope for rapid rescue via new beneficial insertions in a low-recombination genome. Although not addressed here, the polygenic nature of climate adaptation highlights the need for future experimental work to dissect how TE-derived variation collectively shapes adaptive responses.

In summary, our study reveals that ancient transposable element variation can decouple ecological adaptation from nucleotide diversity, offering a general mechanism by which evolution proceeds under persistent genetic paucity.

## Materials and Methods

### Global accessions and sequencing

We analyzed whole-genome sequencing data from a global panel of 245 *Spirodela polyrhiza* accessions (Table S4). Owing to predominantly clonal reproduction, individual accessions are expected to capture local population-level genetic variation. This collection included 228 previously published resequencing datasets (29 from Canada and the USA ^23^, 68 and 131 accessions sampled globally ^25,53^) and 17 newly sequenced accessions generated in this study to improve representation from under-sampled regions of North America.

These 17 additional accessions were collected along a latitudinal gradient across the USA during 2022 and 2023, including populations from New York, Ohio, Indiana, Missouri, Mississippi, and Texas. Following established protocols ^29^, axenic stocks of accessions were established and cultured in 0.5× Hoagland medium (PhytoTech Labs, Lenexa, Kansas) supplemented with 0.5% sucrose at 24°C under a 16-h light/8-h dark photoperiod. For each accession, approximately 60 clonal individuals were silica-dried and used for genomic DNA extraction using the E.Z.N.A. SP Plant DNA Kit (Omega Bio-Tek Inc., Norcross, Georgia), followed by elution in 100 μL of sterile TE buffer. Whole-genome sequencing libraries were prepared and sequenced at Novogene on a NovaSeq X Plus platform (paired-end 150 bp).

The combined dataset was generated on multiple Illumina platforms: HiSeq2000 (PE 100 bp; mean depth of ∼17×) ^23^, HiSeq X Ten (PE 150 bp; ∼45×) ^53^, HiSeq 4000 (PE 150 bp; ∼38×) ^25^, and NovaSeq X Plus (PE 150 bp; *∼*120×, this study).

### SNP calling and filtering

Chromosome-level genome assemblies are currently available for two *Spirodela polyrhiza* clones (7498 ^54^ and 9509 ^43,55^). We selected the most recent assembly of *Spirodela polyrhiza* 9509 ^55^ as the reference genome due to its superior contiguity (N50 = 7.95 Mb) and improved structural and functional annotation.

Raw reads from the combined global dataset were quality filtered using TRIMMOMATIC v.0.39 ^56^ to remove adapters, potential contamination, and low-quality bases, with the following parameters: LEADING:3 TRAILING:3 SLIDINGWINDOW:4:15 MINLEN:36. Clean reads were aligned to the reference genome using BWA-MEM ^57^ with default parameters. To minimize potential biases from sequencing depth variation among accessions, all BAM files were down-sampled to a maximum depth of 30× using SAMtools v1.9 with the -s parameter ^58^. PCR duplicates were removed using the rmdup function in SAMtools.

SNP calling was performed using the Genome Analysis Toolkit (GATK) v4.3.0.0 ^59^. Per-sample GVCFs were generated using HaplotypeCaller, merged with CombineGVCFs and jointly genotyped using GenotypeGVCFs. Variants were filtered using VariantFiltration with the following thresholds: QD < 3.0, FS > 80.0, SOR > 4.0, MQRankSum < −12.5 and ReadPosRankSum < −8.0. Additional filtering removed non-biallelic sites using BCFtools v1.6 ^60^, variants with >20% missing data using VCFtools v0.1.17 ^61^ (--max-missing 0.8), and SNPs located in organelle genomes, yielding 3,180,216 high quality SNPs. Linkage disequilibrium (LD) pruning was performed in PLINK v1.90b6.21 ^62^ (--indep-pairwise 100 10 0.95), resulting in 1,675,572 independent SNPs. Variants were annotated using SnpEff ^63^, and four-fold degenerate transversion (4DTv) sites were extracted as putatively neutral SNPs (*n* = 146,267) for population structure and demographic inference.

### Population structure

To infer population structure, we retained unrelated accessions from the global collection. Pairwise relatedness was estimated using NgsRelate ^64^, and individuals with rab > 0.95 were grouped into clonal families, resulting in 183 families from the 245 accessions. For each family, the accession with the highest sequencing depth and available geographic coordinates was retained.

Population structure was inferred using the 146,267 4DTv sites from the 183 representative accessions. Principal component analysis (PCA) was performed using PLINK. Ancestry proportions were estimated using ADMIXTURE v.1.3.0 ^65^ with *K* values ranging from 1 to 10 and cross-validation (--cv -j10).

To further characterize genome-wide genetic relationships, maximum likelihood (ML) trees were reconstructed from the same 4DTv dataset using IQ-TREE v2.4.0 ^66^ (-b 500) and RAxML-NG v. 1.2.2 ^67^ (--model GTR+G --all --bs-trees 100). Two accessions of *Colocasia esculenta*, a relative within the same family (Araceae) as *Spirodela*, were included as outgroups. Raw Illumina reads of *Colocasia esculenta* (NCBI: DRR596644, DRR596643) were processed using the same quality control, alignment and SNP calling pipeline as described above, with reads aligned to the *Spirodela polyrhiza* 9509 reference genome. As IQ-TREE and RAxML produced congruent topologies, the result from IQ-TREE was presented.

### TreeMix analysis

To infer population relationships and historical migration events, we applied TreeMix v1.13 ^27^ to the same 4DTv dataset of the 183 representative accessions. Although the 4DTv sites had previously been pruned for LD, additional LD pruning was performed using PLINK (--indep-pairwise 100 20 0.2) according to TreeMix requirements. Variants were further filtered to remove missing data using VCFtools (--max-missing 1), leading to 74,012 SNPs. TreeMix was run with the number of migration edges (m) ranging from 0 to 15, using -k 300 to account for residual LD and -bootstrap 100 to assess node support. The optimal model was determined by examining changes in log-likelihoods across successive values of m and identifying where additional migration edges resulted in minimal improvement.

### Demographic history inference

Demographic history was inferred using complementary coalescent approaches. Multiple Sequentially Markovian Coalescent 2 (MSMC2 v2.1.4) ^68^ was used to infer changes in effective population size (*N*e) through time. To ensure robustness of inference, especially for recent *N*e dynamics ^69^, we additionally applied Sequentially Markovian Coalescent method (SMC++) ^70^. To test alternative demographic scenarios and estimate divergence times, migration rates and *N*e, we further applied fastsimcoal2 v.2.7.0.9 ^71^ using the joint site frequency spectrum.

For *N*e inference using MSMC2 and SMC++, both variant and invariant sites were required. Therefore, an all-sites dataset was generated by re-running joint genotyping with GenotypeGVCFs using the “-all-sites” option. This yielded 122,156,028 callable sites across the 183 representative individuals, including 3,083,943 variant sites and 119,072,085 invariant sites.

As MSMC2 infers coalescence events between haplotypes separated by recombination, we assume that vegetative propagation does not involve any coalescence events between haplotypes or recombination, and sexual reproduction is the only time that recombination and coalescence occur. We therefore parameterized demographic inference using a sexual generation time and mutation rate. Based on an estimated mutation rate (µ) of 2.38 × 10^−10^ per site per asexual generation (3 days) in *Spirodela polyrhiza* ^53^, we assumed a sexual generation time of 20 days ^72^ (gen = 0.055), and corresponding to a mutation rate of 1.67 × 10^−9^ per site per sexual generation. Although the average time between sexual generations in natural populations is not precisely known, sensitivity analyses showed that varying the assumed sexual generation time (with corresponding mutation rate adjustments) altered absolute estimates of *N*e but did not affect the inferred timescale in years. Accordingly, while absolute *N*e values should be interpreted with caution, the absolute timing of events is robust to our assumption of sexual generation time.

For MSMC2 analysis, two individuals (four haplotypes) were selected to represent each population (inferred based on population structure section above), with samples drawn from different geographic locations. Input files were generated using two kinds of masks: individual-level positive masks based on mapping coverage and a single negative mask corresponding to coding (CDS) regions. Multihetsep files were generated separately for each chromosome using generate_multihetsep.py. The time segment patterning was defined as “-p 1*2+15*1+1*2”. To assess the robustness of *N*e inference, we performed 100 bootstrap replicates at the multihetsep level using multihetsep_bootstrap.py. Briefly, the genome was partitioned into non-overlapping segments of fixed size, and bootstrap datasets were generated by random resampling these genomic segments with replacement across chromosomes. MSMC2 was run independently for each bootstrap replicate using identical parameters. Relative cross-coalescence rates (RCCRs) were estimated between populations (each represented by four haplotypes) using combineCrossCoal.py. The RCCR ranges from 0 to 1, with a value of 0.5 approximating the splitting time between populations.

The inference of *N*e dynamics using SMC++ used parameters “--timepoints 1000 1000000” and the mutation rate of 1.67 × 10^−9^ per site per sexual generation. Confidence intervals were obtained using the “estimate” command in SMC++ and visualized with the “-c” option.

For fastsimcoal2 analysis, we constructed two-dimensional (2D) joint folded site frequency spectra (SFSs) of neutral sites that include both variant and invariant positions. Neutral sites were extracted from the all-sites dataset (122,156,028 callable sites across 183 individuals), retaining 3,000,759 intergenic sites located at least 10 kb from annotated genes to minimize the influence of selection. SFSs were generated using easySFS.py (https://github.com/isaacovercast/easySFS) with parameters “--proj 24, 24, 24, 24”. Guided by lineage relationships inferred from MSMC2 and SMC++, we formulated three alternative demographic models (Fig. S2) that shared the same branching topology but differed in the inclusion and geographic placement of post-divergence bottlenecks. Model A included bottlenecks in North America (NA) and India (IN) immediately following lineage splitting, consistent with dispersal out of Asia. Model B included no post-split bottlenecks, representing a scenario without explicit source of directional dispersal. Model C included a bottleneck in Asia (AS) following lineage splitting, consistent with dispersal out of North America. For each model, 100 independent optimization runs were performed with 100,000 coalescent simulations and 40 cycles of the expectation–conditional maximization (ECM) algorithm. The best-fitting model was selected using Akaike’s information criterion (AIC) and Akaike’s weights. Parameter uncertainty was assessed using 100 parametric bootstrap replicates to derive 95% confidence intervals.

### Genetic diversity

Population-level genetic diversity was quantified using nucleotide diversity (π; the average number of nucleotide differences per site between two randomly chosen sequences within a population), pairwise genetic differentiation (*F*st), and absolute divergence (*d*xy; the average number of nucleotide differences per site between sequences from two different populations).

Estimates were calculated using *pi*xy v.1.2.7.beta1 ^73^ with non-overlapping 50-kb windows. Individual-level genetic diversity was quantified using observed heterozygosity (*H*et) and inbreeding coefficient (*F*) using PLINK. For inbreeding coefficient estimation, allele frequencies were estimated within populations, such that *F* reflects the deviation of observed heterozygosity from Hardy-Weinberg expectations based on population-specific allele frequencies. All estimates were based on the all-sites dataset (122,156,028 callable sites across 183 individuals) including both variant and invariant sites.

### TE polymorphisms and age estimation

To identify TE polymorphisms in the global collection, we first annotated TEs in the *Spirodela polyrhiza* 9509 reference genome using EDTA v2.2.2 ^74^, resulting in 604 TE consensus sequences that were used to construct a species-specific TE library.

TE polymorphisms were detected from the quality-filtered reads using the TEmarker pipeline ^75^. Polymorphisms were inferred based on discordant, split and clipped reads spanning TE–flank junctions ^75^. To increase robustness, four complementary TE variation detection tools wrapped in McClintock ^76^ integrated within TEmarker were used, including ngs_te_mapper2 ^77^, TEMP2 ^78^, PoPoolationTE2 ^79^, and TE-Locate ^80^. Polymorphism calls from these tools were integrated to generate a pan-TE set across all individuals, merging closely located insertion sites into unified TE loci and removing non-polymorphic loci. Then, TE genotypes were assigned based on the proportion of breakpoint-supporting reads at each candidate insertion site. Analyses were conducted with default parameters, except filtering out TE loci using with threshold “-f_th 0.02”. In total, 14,094 polymorphic TEs were identified and were broadly distributed along chromosomes. Functional annotation of polymorphic TEs was performed using ANNOVAR ^81^.

TE insertion ages were estimated following established methods by Baduel et al. (2021) ^11^ and Ren et al. (2025) ^22^. For each polymorphic TE, individuals carrying this insertion were identified using BCFtools v1.6 ^82^, and pairwise SNP differences among carriers were calculated within a 70-kb window centered on the insertion using PLINK. The maximum pairwise SNP difference (*N*_max_) was used to estimate insertion age according to:

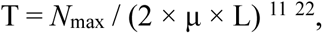

where T denotes insertion time (year), L is the total flanking sequence length examined (70 kb), and μ is the mutation rate per site per year. The mutation rate was set to 3.04 × 10^-^⁸ per site per year, derived from 1.67 × 10^−9^ per site per sexual generation (20 days) in *Spirodela polyrhiza*.

### Genotype-environment association

Nineteen bioclimatic variables representing recent climate conditions (1970 - 2000) were obtained from the WorldClim version 2.1 database (https://worldclim.org/) at 2.5 arc-minute resolution using the R packages geodata ^83^ and terra ^84^. This spatial resolution closely matched the precision of the global collections. To reduce multicollinearity among climatic predictors, PCA was performed using the prcomp() function in R v4.4.1 ^85^. Variables contributing strongly to the first two principal components were retained, while highly correlated predictors were excluded (pairwise Pearson’s |*r*| ≥ 0.8). This procedure yielded six bioclimatic variables: Bio06 (Minimum Temperature of Coldest Month), Bio07 (Temperature Annual Range), Bio12 (Annual Precipitation), Bio15 (Precipitation Seasonality), Bio17 (Precipitation of Driest Quarter), and Bio19 (Precipitation of Coldest Quarter).

Genotype-environment associations (GEAs) were performed using redundancy analysis (RDA) implemented in the R package vegan ^86^ and latent factor mixed models (LFMM) implemented in the R package LEA ^32^. GEAs were conducted for both polymorphic TEs and SNPs across 238 individuals with known geographic coordinates. For SNP-based analyses, variants were filtered for minor allele frequency (MAF ≥ 0.05) from the set of 1,675,572 high-quality, LD-pruned SNPs using PLINK, resulting in 232,169 SNPs retained for GEAs.

For RDA-based GEAs, partial RDA was conducted to control for population structure and spatial autocorrelation by conditioning on the first two principal components of genome-wide variation as well as geographic coordinates (latitude and longitude). Individual-level genotypes were used as the response matrix, with the six selected bioclimatic variables as predictors. The relative contributions of bioclimatic variables were assessed using 1000 permutations (anova, by = “margin”). Climate-associated loci were identified using the rdadapt framework ^31^, based on SNP or TE loadings along constrained RDA axes (up to five axes). To reduce false positives, null models were generated by randomizing geographic coordinates (i.e., environmental variables) among individuals, thereby disrupting genotype-environment associations. The distribution of *P* values from 10 randomized runs was compared with that of the observed data to determine an empirical threshold (20% here) above which associations exceeded random expectation ^87^.

For LFMM-based GEAs, population structure was accounted for by estimating ancestry coefficients using sNMF implemented in the R package LEA. The number of ancestral populations (*K*) was evaluated from *K* = 2 to *K* = 10 using the cross-entropy criterion, with *K* = 7 showing the lowest cross-entropy (consistent with population structure inferred above). This value was therefore used as the number of latent factors in LFMM. SNP and TE genotype matrices were converted to LFMM format using ped2lfmm(). Missing genotypes were imputed via impute() in LEA, and SNP genotype imputation was carried out using Beagle v5.4 ^88^ to improve computational efficiency. LFMM was run with five repetitions, 2,000 iterations per run, and a burn-in of 500 iterations. Association *P* values were obtained with lfmm.pvalues() and adjusted for multiple testing using Bonferroni correction. As in RDA analysis, null models were generated by randomly permuting geographic coordinates among individuals, and the full analysis was repeated 10 times to derive an empirical threshold for identifying climate-associated loci.

### Climate adaptive loci

While GEAs identify loci correlated with environmental gradients, such associations may arise from processes other than selection. To more robustly infer climate adaptation, we therefore identified candidate climate adaptive loci by integrating evidence from both GEAs and independent signatures of selection.

To identify loci under selection independent of environmental association, we applied an ancestry-based outlier detection approach based on sNMF implemented in LEA. This method detects loci exhibiting stronger differentiation than expected under the genome-wide admixture model (*K* = 7). Loci with unusually large deviations from neutral population structure were considered candidates for selection. To reduce false positives, we generated an empirical null distribution of *P* values using the putatively neutral 4DTv dataset (146,267 SNPs). SNP and TE loci with observed *P* values falling below the 5th percentile of the null distribution (empirical *P* < 0.05) were retained as candidate loci under selection. Climate adaptive loci were defined as the intersection between loci showing significant genotype–environment associations and those identified as candidates under selection.

To ensure the robustness of inference for the identified climate adaptive loci, we further evaluated whether these loci were indeed under selection and assessed the type of selection (positive, balancing or purifying selection) using BayeScan v2.1 ^89^. BayeScan was applied to the set of SNP and TE loci included in GEAs, with VCF files converted into BayeScan-compatible input format using PLINK. Each polymorphic TE was treated as a biallelic locus, analogous to SNPs. The Markov Chain Monte Carlo (MCMC) algorithm was run with a burn-in of 50,000 iterations, followed by 5,000 output iterations sampled every 10 iterations (thinning interval = 10). Twenty pilot runs of 5,000 iterations were conducted to calibrate the proposal distributions. The prior odds for the neutral model were set to 10. Loci under selection were identified based on posterior probabilities and associated q-values (< 0.05). For loci identified as climate adaptive, the sign of the BayeScan α parameter was used to infer the direction of selection, with positive α indicating positive (diversifying) selection and negative α indicating balancing or purifying selection.

### Genetic offset of climate adaptive loci

To assess future climate–genotype mismatch, we estimated genetic offset using two complementary approaches: a site-scale LFMM-based approach calculated with LEA and a range-scale genetic offset calculated with the R package gradientForest ^40^. Because relatively few SNPs were identified as climate adaptive loci, analyses focused on climate adaptive TEs and the strongest climate selective axis identified (Minimum Temperature of Coldest Month, Bio6).

For the site-scale analysis, genetic offset was calculated using the genetic.gap() function in LEA, with latent factors (*K* = 7) included to account for population structure. For the range-scale analysis, gradient forest models with 500 regression trees were used to model the relationship between TE genotypes and Bio6. To further account for population structure, scores along the first principal component (PC1) were included as a covariate. Future climate projections (2081-2100, SSP5-8.5 scenario) were used to estimate genetic offset. Differences between populations were assessed using two-sided Wilcoxon tests with Benjamini-Hochberg correction for multiple comparisons.

### Genomic context of climate adaptive TEs

To examine the genomic context of climate adaptive TEs, the genome was partitioned into fixed, non-overlapping 10-kb windows. A window was defined as a climate adaptive TE window if it harbored at least one climate adaptive TE. Windows that contained non-climate-adaptive TEs were defined as non-adaptive windows and used as a reference set for comparison. We compared π, Tajima’s D, and selective sweep signals between climate adaptive TE windows and non-adaptive TE windows using the full set of high-quality SNPs (3,180,216 SNPs across 183 representative accessions). π and Tajima’s D were estimated using VCFtools. Selective sweep signals were assessed using SweeD v3.0 ^38^, and the composite likelihood ratio (CLR) was estimated based on deviations from the neutral site frequency spectrum and averaged within corresponding windows.

Because non-adaptive windows greatly outnumbered adaptive TE windows, we randomly subsampled non-adaptive windows to match the number of adaptive windows 1,000 times, generating a permutation-based null distribution for each statistic (π, Tajima’s D, and mean CLR). To test whether statistics calculated for adaptive windows differed significantly from non-adaptive windows (null expectation here), we computed a two-sided permutation *P* value as:

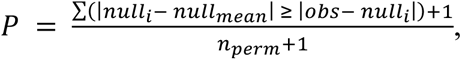

where *obs* denotes the observed statistic across adaptive windows, *null*_i_ represents individual value from each permutation of non-adaptive windows, *null*_mean_ is the mean of the null distribution, and *n*_perm_ = 1000 is the number of permutations. This formulation quantifies the proportion of permuted values that are at least as extreme as the observed statistic relative to the null expectation.

### Ages of climate adaptive vs. non-adaptive TEs

To test whether climate adaptive TEs differed in age from non-adaptive TEs, we compared the estimated insertion ages between these two groups of polymorphic TEs. Because non-adaptive TEs outnumbered adaptive TEs, we randomly subsampled non-adaptive TEs to match the number of adaptive TEs 1,000 times. Statistical significance was assessed using a two-sided permutation test as described above.

### π_N_/π_S_ estimation

To quantify variation in selective constraint across the genome, we estimated the ratio of nonsynonymous to synonymous nucleotide diversity (πN/πS) in non-overlapping 10-kb windows using the full set of high-quality SNPs (3,180,216) across 183 representative accessions. SNPs were annotated with SnpEff; *synonymous* variants were classified as synonymous, and *missense*, *nonsense*, *stop_gained*, or *stop_lost* were classified as nonsynonymous. Within each genomic window, πS and πN were calculated as the mean π across synonymous and nonsynonymous SNPs, respectively. Windows lacking informative variants were excluded, and only windows containing at least 10 SNPs were retained for analysis.

We further compared πN/πS between windows containing TEs (TE windows) and those lacking TEs (non-TE windows). Within TE windows, πN/πS was additionally compared between windows harboring climate adaptive TEs and those containing non-adaptive TEs. Because the number of windows differed substantially between comparison groups, we performed subsampling (1,000 permutations) and assessed statistical significance using a two-sided permutation test as described above.

### Gene Ontology analysis

Functional annotation and Gene Oncology (GO) terms of *Spirodela polyrhiza* 9509 reference proteins were obtained from https://www.lemna.org/ ^55^. GO enrichment analysis was performed using the R package ClusterProfiler ^90^. Enrichment significance was assessed using Fisher’s exact test, and *P* values were adjusted for multiple testing using the Benjamini-Hochberg correction. GO terms with an adjusted *P* value < 0.05 were considered significantly enriched.

## Acknowledgements

This work was supported by the National Science Foundation of China (32571740 to N.W.); Natural Science Foundation of Hubei Province, China (JCZRQNA202600231 to N.W.); Chinese Academy of Sciences (E529990101 and E455990101 to N.W.). We thank Maris Hollowell for assistance with field collection and laboratory work and the Wei Lab members for helpful discussions on visualization.

## Author contributions

N.W. designed the study. A.Z. conducted the data analyses and prepared the figures. N.W. and A.Z. drafted the manuscript. J.B.B. provided guidance on data analyses, and all authors contributed to manuscript revision.

## Supplementary information

The online version contains supplementary material.

## Data Availability

Whole genome sequencing data used in this study were obtained from the NCBI Sequence Read Archive under accession numbers SRP213917, SRP306079, SRP154628. Newly generated sequencing data and other source data from this study have been deposited in SciDB (https://www.scidb.cn/) and will be publicly available upon publication.

