## Supporting Information for "Ancient transposable elements sustain global ecological adaptation despite chronically low nucleotide diversity"

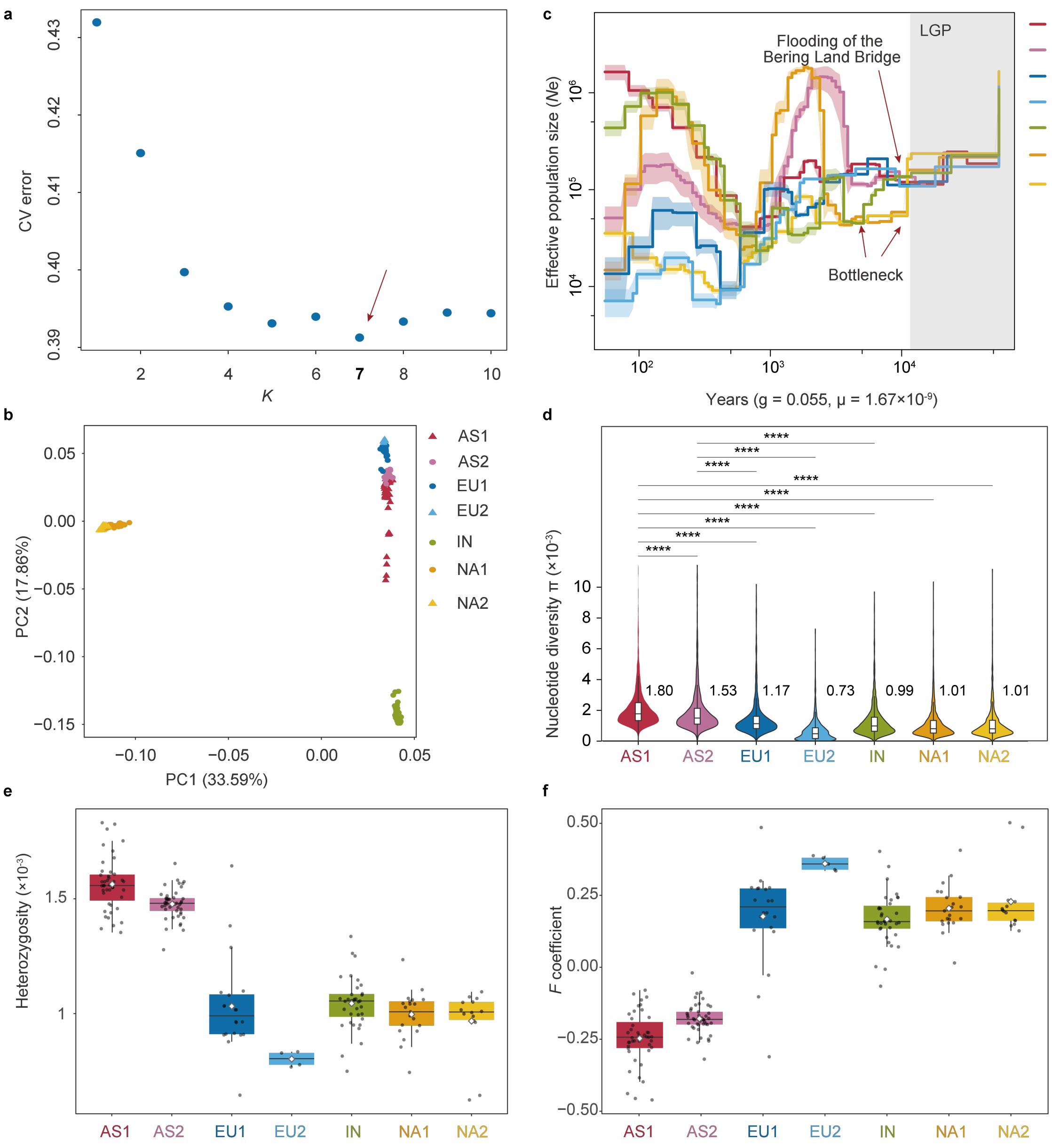


**Fig. S1. Genetic structure and summary of population genetic parameters of *Spirodela polyrhiza*.** (a) The optimal number of genetic clusters (*K)* value inferred from cross-validation analysis in ADMIXTURE v.1.3.0 [1] with parameters "--cv -j10", based on 146,267 putatively neutral 4DTv sites. (b) Principal Component analysis (PCA) based on unrelated neutral SNPs from 183 individuals with PLINK v1.90b6.21 [2]. Variance explained by PC1 and PC2 is shown in parentheses. (c) The dynamics of effective population size (Ne) using SMC++ [3], Estimates assumed a sexual generation of 20 days (gen = 0.055) and a mutation rate of 1.67 × 10-9 per site per sexual generation. Major demographic events are indicated, including colonizationassociated bottlenecks and the final inundation of the Bering Land Bridge. (d) Comparison of nucleotide diversity ($\pi$) based on SNPs. (e, f) Distribution of individual heterozygosity and inbreeding coefficient ($F$). The white boxes in the violin plots represent the interquartile range, and the black lines indicate the medians. Pairwise differences were assessed using two-sided Wilcoxon rank-sum tests (*** $P < 0.001$; ** $P < 0.01$, * $P < 0.05$).

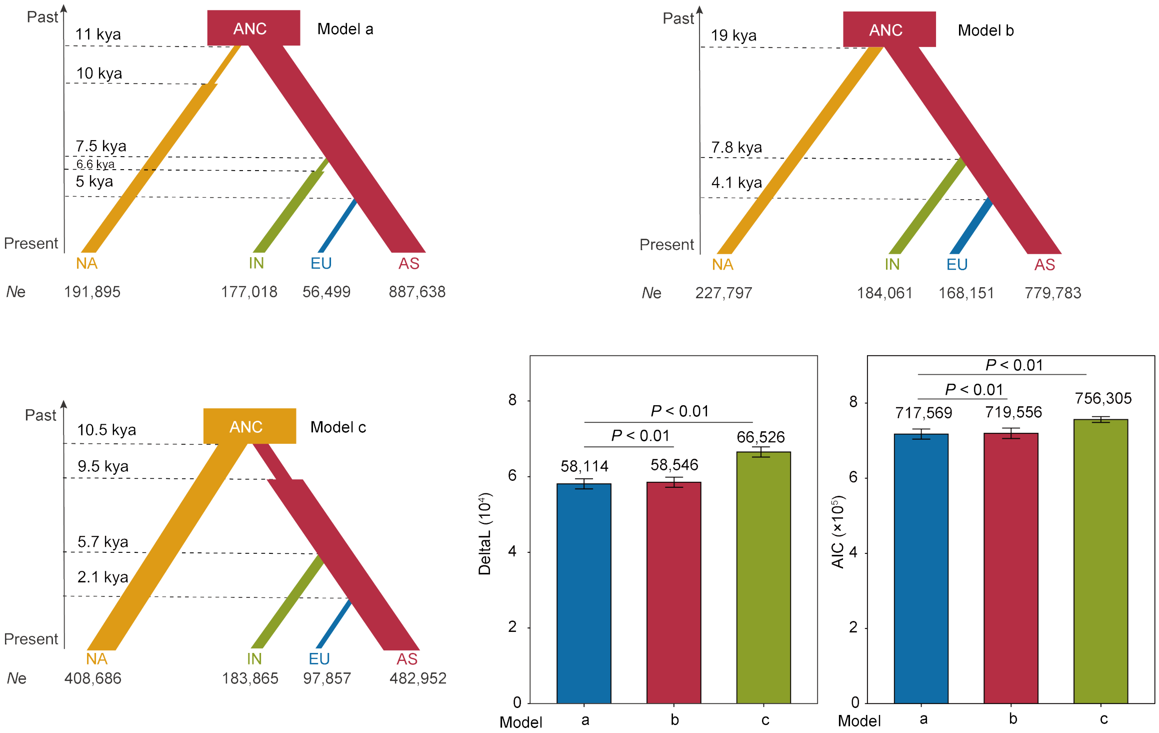


**Fig. S2. Schematic diagram of 3 models with different demographic history and bottleneck occurrence used for fastsimcoal2 [4] simulation of *Spirodela polyrhiza*.** All models shared the same branching topology but differed in whether post-dispersal bottlenecks were included. Model a incorporated bottlenecks after lineage splitting in NA and IN, consistent with dispersal out of Asia. Model b included no post-split bottlenecks, representing an unresolved ancestral source. Model c incorporated bottlenecks after lineage splitting in AS, consistent with dispersal out of North America. Boxes represent lineages, with widths proportional to estimated effective population sizes, 'ANC' represents the common ancestor of four main populations. Gray arrows denote gene flow (per generation migration rate). Divergence times are indicated as kya, the estimates of *N*e are shown below. Model a was selected as the best model according to the lowest DeltaL and Akaike information criterion (AIC) values. Significance of pairwise comparisons was assessed using the two-sided Wilcoxon rank-sum tests.

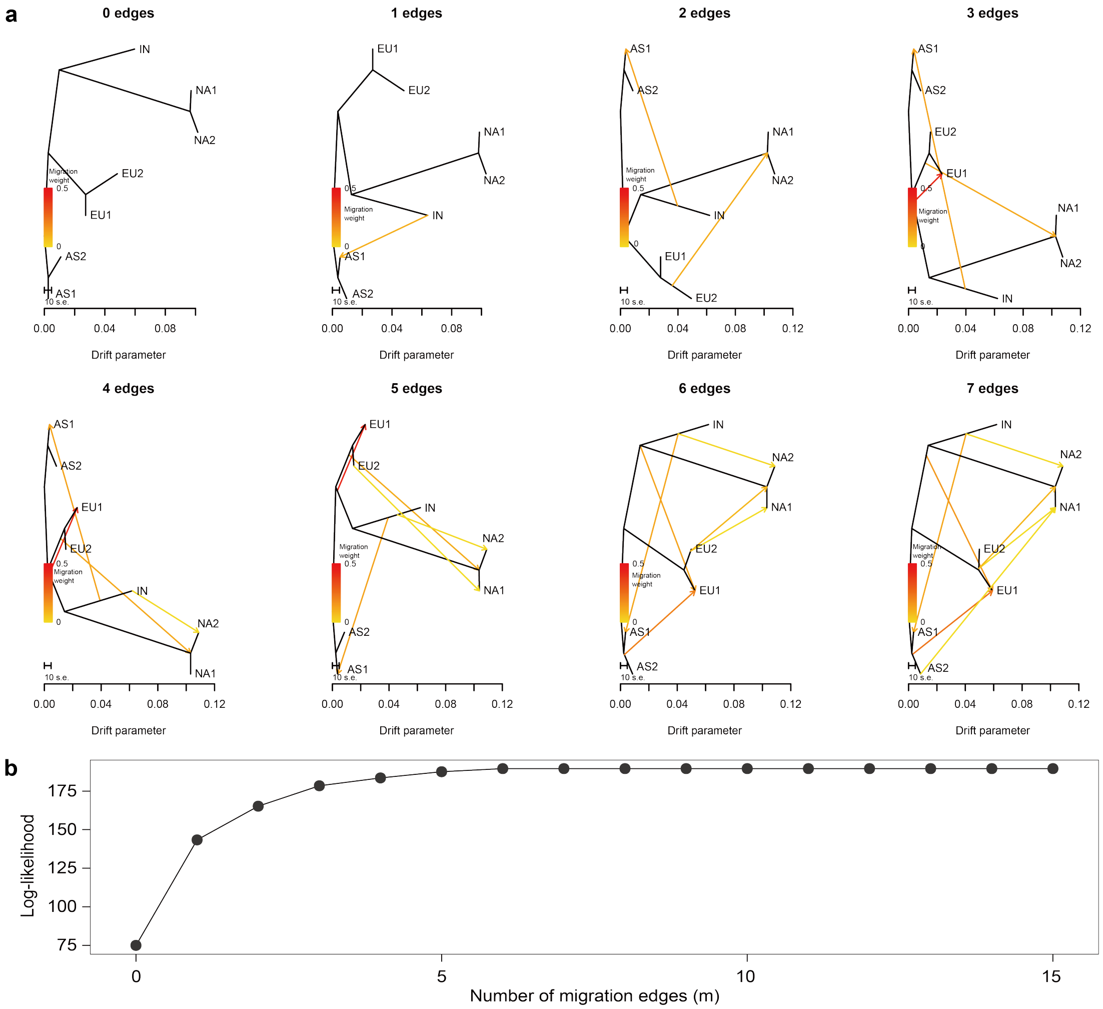


**Fig. S3. Gene flow events inferred by Treemix of *Spirodela polyrhiza*.** (a) Gene flow inferred from TreeMix v1.13 [5] with increasing numbers of migration edges (m = 0 - 7). Arrows denote migration events, and color intensity represents migration weight. (b) Model selection for TreeMix based on maximum likelihood across migration edges. The optimal model was determined by examining changes in log-likelihoods across successive values of m and identifying where additional migration edges resulted in minimal improvement. Here, m = 6 was selected as the optimal number of migration edges.

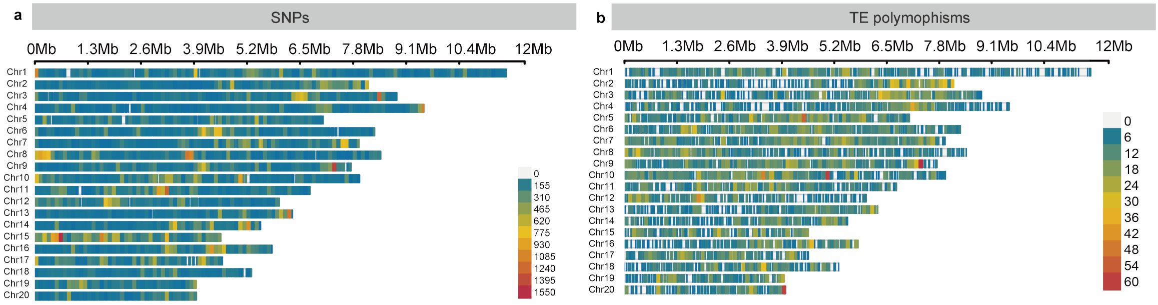


**Fig. S4. Chromosomal distribution of genomic variation of *Spirodela polyrhiza.*** (a) Chromosomal distribution of single-nucleotide polymorphism (SNPs). (b) Chromosomal distribution of transposon elements (TE) polymorphisms. The blocks are colored by distribution density calculated in 100-kb windows across the genome, with red indicating high and blue indicating low.

a

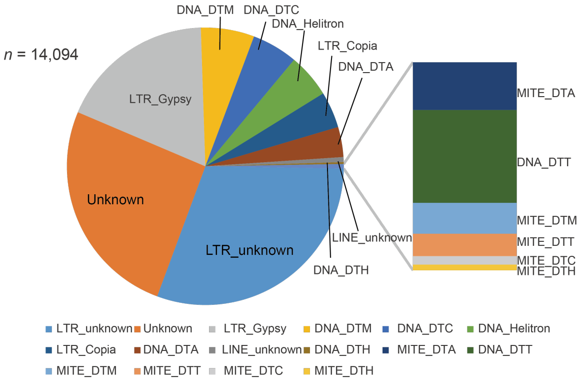


| Type | Common Name | Reference | TE polymorphism |
|---|---|---|---|
| DNA_DTA | hAT DNA | 2,561 | 494 |
| DNA_DTC | CACTA DNA | 4,139 | 754 |
| DNA_DTH | PIF/Harbinger DNA | 192 | 31 |
| DNA_DTM | Mutator DNA | 4,633 | 880 |
| DNA_DTT | Tc1/Mariner DNA | 149 | 33 |
| DNA_Helitron | Helitron DNA | 3,684 | 723 |
| LINE_unknown | Long Interspersed Elemen | 307 | 82 |
| LTR_Copia | Ty1/Copia | 3,163 | 596 |
| LTR_Gypsy | Ty3/Gypsy | 9,180 | 2,551 |
| LTR_unknown | Unknown LTR | 19,751 | 4,285 |
| MITE_DTA | hAT-derived MITE | 162 | 17 |
| MITE_DTC | CACTA-derived MITE | 32 | 3 |
| MITE_DTH | Harbinger-derived MITE | 30 | 2 |
| MITE_DTM | Mutator-derived MITE | 78 | 11 |
| MITE_DTT | Tc1/Mariner-derived MITE | 34 | 8 |
| Unknown | Unknown | 17,439 | 3,624 |

b c

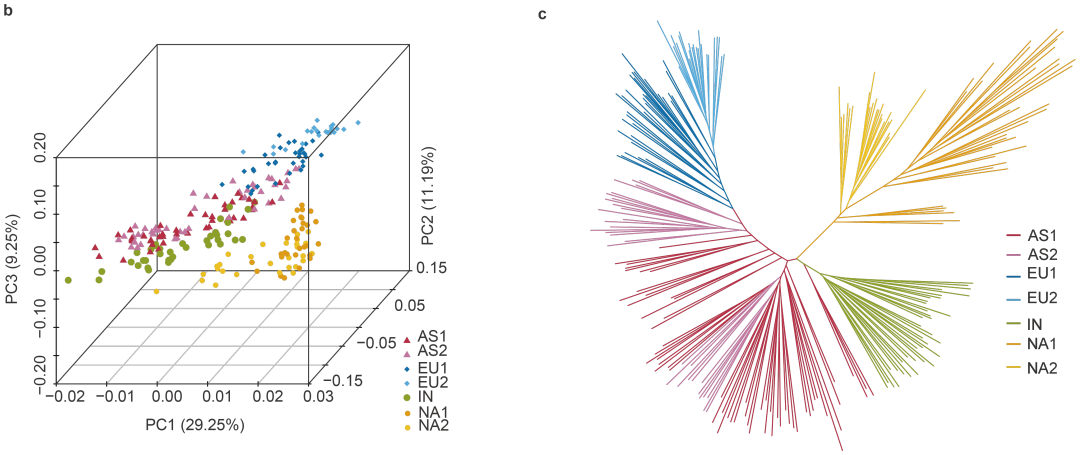


**Fig. S5. Characteristics of TE polymorphisms of *Spirodela polyrhiza*.** (a) Distribution of major TE superfamilies in the reference genome and TE polymorphisms detected across populations of *Spirodela polyrhiza*. (b) Principal component analysis (PCA) based on TE polymorphisms, revealing the distinct genetic structure. The percentage of variation explained by the first three PCs are shown in parentheses. (c) Maximum likelihood (ML) phylogeny inferred from genome-wide TE polymorphisms. The clades are colored by lineages based on SNPs.

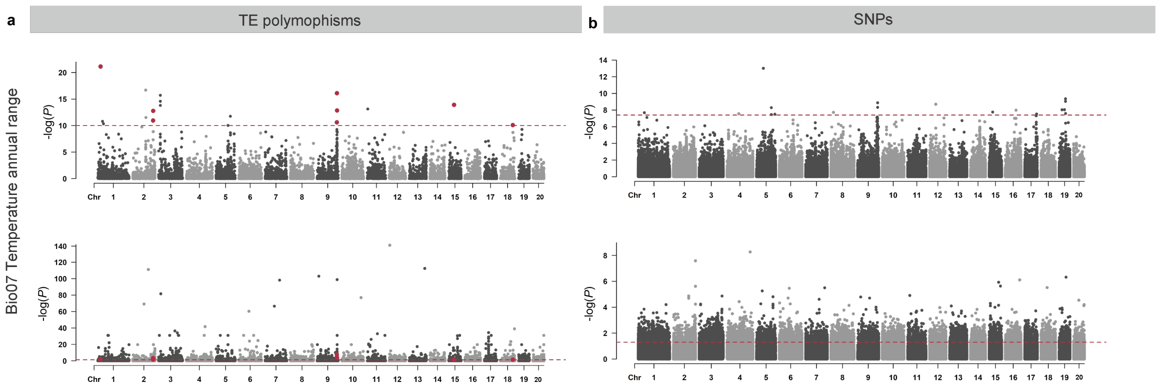


**Fig. S6. Manhattan plots of climate adaptive TE polymorphisms and SNPs associated with the annual temperature range (Bio07).** Climate adaptive TE polymorphisms and SNPs are shown in (a) and (b), respectively. Loci supported by both GEAs (LFMM; upper panels) and selection signals inferred by sNMF (lower panels) were defined as climate adaptive. Dashed red lines in the upper panels indicate an empirical threshold derived from null models generated by randomizing geographic coordinates to disrupt genotype-environment associations, whereas dashed red lines in the lower panels indicate the empirical threshold derived from putatively neutral 4DTv sites.

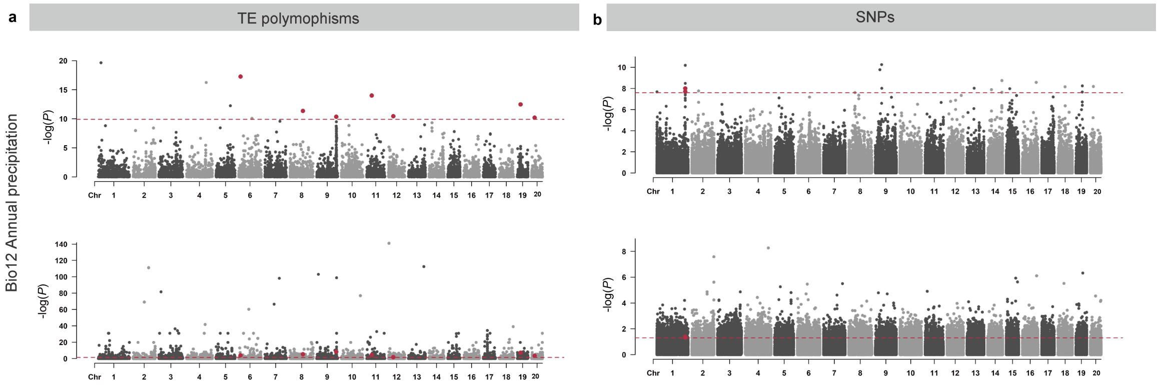


**Fig. S7. Manhattan plots of climate adaptive TE polymorphisms and SNPs associated with the annual precipitation (Bio12).** Climate adaptive TE polymorphisms and SNPs are shown in (a) and (b), respectively. Loci supported by both GEAs (LFMM; upper panels) and selection signals inferred by sNMF (lower panels) were defined as climate adaptive. Dashed red lines in the upper panels indicate an empirical threshold derived from null models generated by randomizing geographic coordinates to disrupt genotype-environment associations, whereas dashed red lines in the lower panels indicate the empirical threshold derived from putatively neutral 4DTv sites.

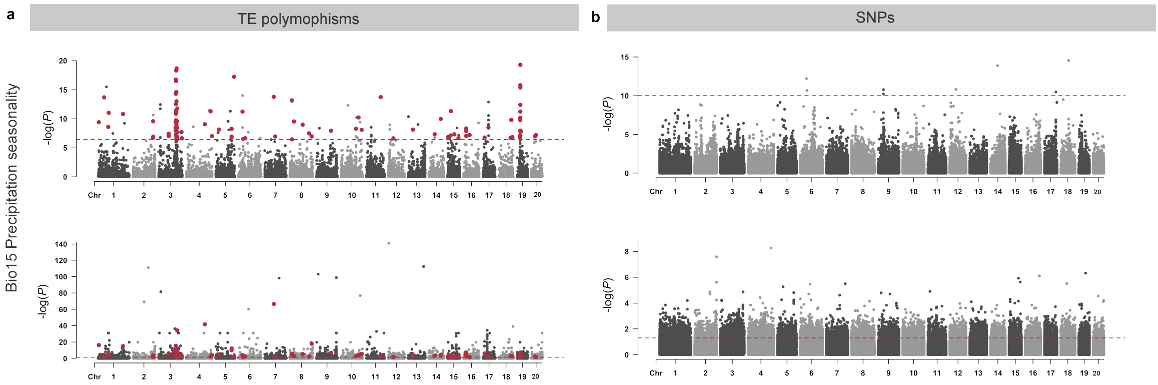


**Fig. S8. Manhattan plots of climate adaptive TE polymorphisms and SNPs associated with the precipitation seasonality (Bio15).** Climate adaptive TE polymorphisms and SNPs are shown in (a) and (b), respectively. Loci supported by both GEAs (LFMM; upper panels) and selection signals inferred by sNMF (lower panels) were defined as climate adaptive. Dashed red lines in the upper panels indicate an empirical threshold derived from null models generated by randomizing geographic coordinates to disrupt genotype-environment associations, whereas dashed red lines in the lower panels indicate the empirical threshold derived from putatively neutral 4DTv sites.

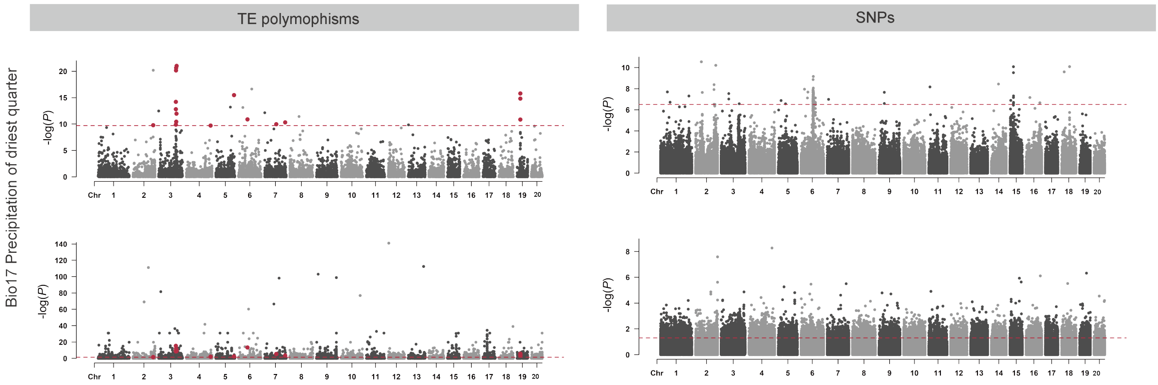


**Fig. S9. Manhattan plots of climate adaptive TE polymorphisms and SNPs associated with the annual precipitation (Bio17).** Climate adaptive TE polymorphisms and SNPs are shown in (a) and (b), respectively. Loci supported by both GEAs (LFMM; upper panels) and selection signals inferred by sNMF (lower panels) were defined as climate adaptive. Dashed red lines in the upper panels indicate an empirical threshold derived from null models generated by randomizing geographic coordinates to disrupt genotype-environment associations, whereas dashed red lines in the lower panels indicate the empirical threshold derived from putatively neutral 4DTv sites.

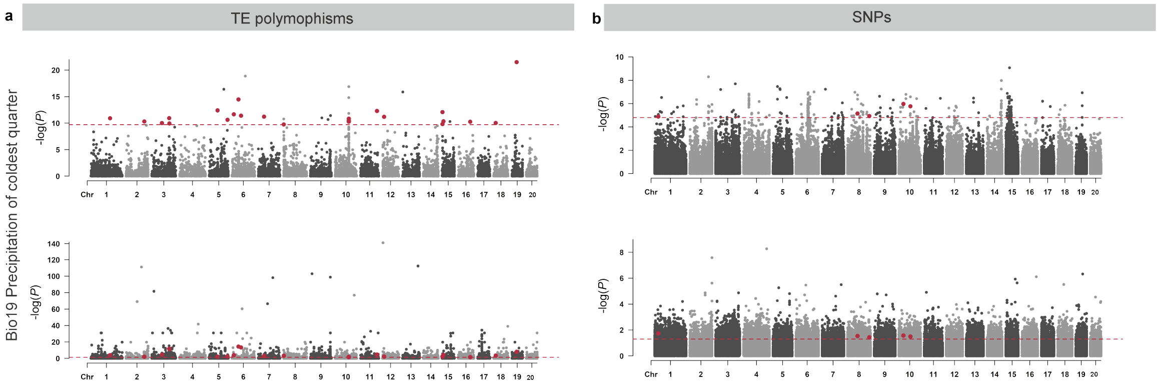


**Fig. S10. Manhattan plots of climate adaptive TE polymorphisms and SNPs associated with the precipitation of the coldest quarter (Bio19).** Climate adaptive TE polymorphisms and SNPs are shown in (a) and (b), respectively. Loci supported by both GEAs (LFMM; upper panels) and selection signals inferred by sNMF (lower panels) were defined as climate adaptive. Dashed red lines in the upper panels indicate an empirical threshold derived from null models generated by randomizing geographic coordinates to disrupt genotype-environment associations, whereas dashed red lines in the lower panels indicate the empirical threshold derived from putatively neutral 4DTv sites.

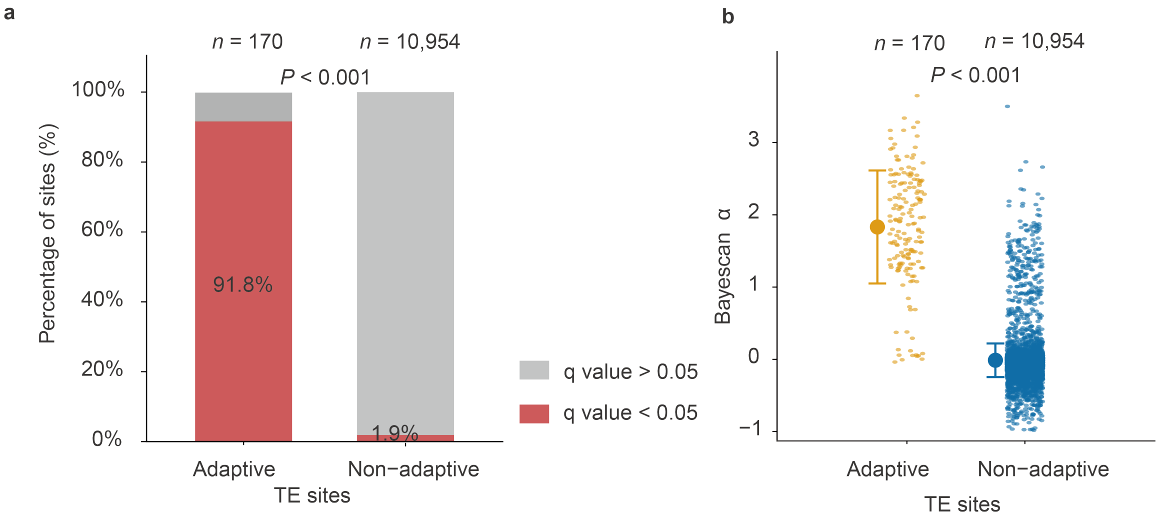


**Fig. S11. Enrichment of selection signals at climate adaptive TE polymorphisms inferred by Bayescan [6].** (a) Proportion of TE polymorphic sites showing significant selection signals (q value < 0.05) among climate adaptive and non-adaptive TE loci. Colors denote loci with significant (red) and non-significant (grey) Bayescan signals. Statistical significance was assessed using a two-proportion z-test. (b) Distribution of Bayescan α values for climate adaptive and non-adaptive TE polymorphisms. Positive α values indicate positive (diversifying) selection, whereas values close to zero or negative indicate balancing or purifying selection, respectively. Because Non-adaptive TEs outnumbered adaptive, non-adaptive TEs were randomly subsampled 1,000 times to generate a permutation-based null distribution. Statistical significance was assessed using a two-sided permutation test.

**Table S1.** Nucleotide diversity (π) estimated from the all-sites dataset (122,156,028 callable sites across 183 individuals) including both variant and invariant sites in *Spirodela polyrhiza*

| Population | Mean | Median | SD | Min | Max |
|---|---|---|---|---|---|
| AS1 | 0.00223 | 0.001801 | 0.001432 | 6.36E-06 | 0.011449 |
| AS2 | 0.001911 | 0.001527 | 0.001331 | 6.70E-06 | 0.011448 |
| EU1 | 0.001456 | 0.001134 | 0.00117 | 2.50E-07 | 0.010184 |
| EU2 | 0.000658 | 0.000483 | 0.000731 | 1.91E-05 | 0.007287 |
| IN | 0.001303 | 0.000987 | 0.001097 | 5.20E-06 | 0.009694 |
| NA1 | 0.001129 | 0.000808 | 0.001015 | 9.50E-06 | 0.010347 |
| NA2 | 0.001122 | 0.000802 | 0.00101 | 2.21E-06 | 0.011164 |

**Table S2.** Distribution of SNP and TEs variant types in *Spirodela polyrhiza*

| Category | Type | Count | Percentage (%) |
|---|---|---|---|
| SNP | exonic | 300,449 | 9.45 |
| | intronic | 681,440 | 21.43 |
| | splicing | 2,305 | 0.07 |
| | upstream;downstream | 379,444 | 11.93 |
| | UTR | 112,852 | 3.55 |
| | intergenic | 1,636,220 | 51.45 |
| TE | exonic | 4,613 | 32.73 |
| | intronic | 1,211 | 8.59 |
| | splicing | 15 | 0.11 |
| | upstream;downstream | 1,507 | 10.69 |
| | UTR | 405 | 2.87 |
| | intergenic | 6,130 | 43.49 |
